# The Himalayan Uplift and the Evolution of Aquatic Biodiversity Across Asia: Snowtrout (Cyprininae: *Schizothora*x) as a Test Case

**DOI:** 10.1101/2020.10.12.336149

**Authors:** Binod Regmi, Marlis R. Douglas, David R. Edds, Karma Wangchuk, Chang Lu, Gopal Prasad Khanal, Pema Norbu, Sangay Norbu, Sonam Dorji, Singye Tshering, Zachary Angel, Tyler K. Chafin, Zachery D. Zbinden, Michael E. Douglas

## Abstract

The Himalayan uplift, a tectonic event of global importance, seemingly disseminated aquatic biodiversity broadly across Asia. But surprisingly, this hypothesis has yet to be tested. We do so herein by sequencing 1,140 base-pair of mtDNA *cytochrome-b* for 72 tetraploid Nepalese/Bhutanese Snowtrout (*Schizothorax spp.*), combining those data with 67 GENBANK^®^ sequences (59 ingroup/8 outgroup), then reconstructing phylogenetic relationships using maximum likelihood/ Bayesian analyses. Results indicate Snowtrout originated in Central Asia, dispersed across the Qinghai-Tibetan Plateau (QTP), then into Bhutan via south-flowing tributaries of the east-flowing Yarlung-Tsangpo River (YLTR). The headwaters of five large Asian rivers provided dispersal corridors into southeast Asia. South of the Himalaya, the YLTR transitions into a westward-flowing Brahmaputra River that facilitated successive colonization of Himalayan drainages: First Bhutan, then Nepal, followed by far-western drainages subsequently captured by the Indus River. We found greater species-divergences across rather than within-basins, implicating vicariant evolution as a driver. The Himalaya is a component of the “third-pole” [Earth’s largest (but rapidly shrinking) glacial reservoir outside the Arctic/Antarctic]. Its unique aquatic biodiversity must not only be recognized (as herein) but also conserved through broad, trans-national collaborations. Our results effectively contrast phylogeography with taxonomy as a necessary first step in this process.

The Himalaya is the most extensive and recently evolved mountain system on Earth (length=2400km; width=240km; elevation=75-8800m), with a global significance underscored by its large-scale lithospheric, cryospheric, and atmospheric interactions [1]. These have not only driven global climate, but also defined the cultural and biological endemism of the region [2]. Massive, tectonically derived mountain chains such as the Alps and the Himalaya are hypothesized as being fundamental to the formation of global biodiversity gradients via vicariance and local adaptation, with a significantly stronger signal in terrestrial rather than aquatic systems [3]. Here we test how orogeny (the deformation and folding of Earth’s crust by lateral compression) has contributed to the diversification of freshwater fishes broadly across Asia. We do so by evaluating the phylogeography of an endemic high-elevation fish, the Snowtrout (*Schizothorax*: Cyprinidae).

## Himalayan orogeny

The Himalaya formed as a series of parallel ranges bordered by rivers on the west (Indus) and east (Brahmaputra), and encompassing most of northern Pakistan, northern India (Kashmir to the west, Assam to the east), as well as Nepal, Bhutan, and parts of China, with the extensive Qinghai-Tibetan Plateau (QTP) to the north and the alluvial plains of India and Bangladesh to the south (Fig. 1). It evolved over a 30my span, the result of a two-stage impact of the Indian craton with the Eurasian landmass. The leading edge of the Indian craton was subducted, and its upper surface essentially ‘skimmed’ so as to form three sequential north-to-south stratigraphic zones (i.e., Tibetan, Greater Himalaya, and Lesser Himalaya [4]).

**Figure 1:**
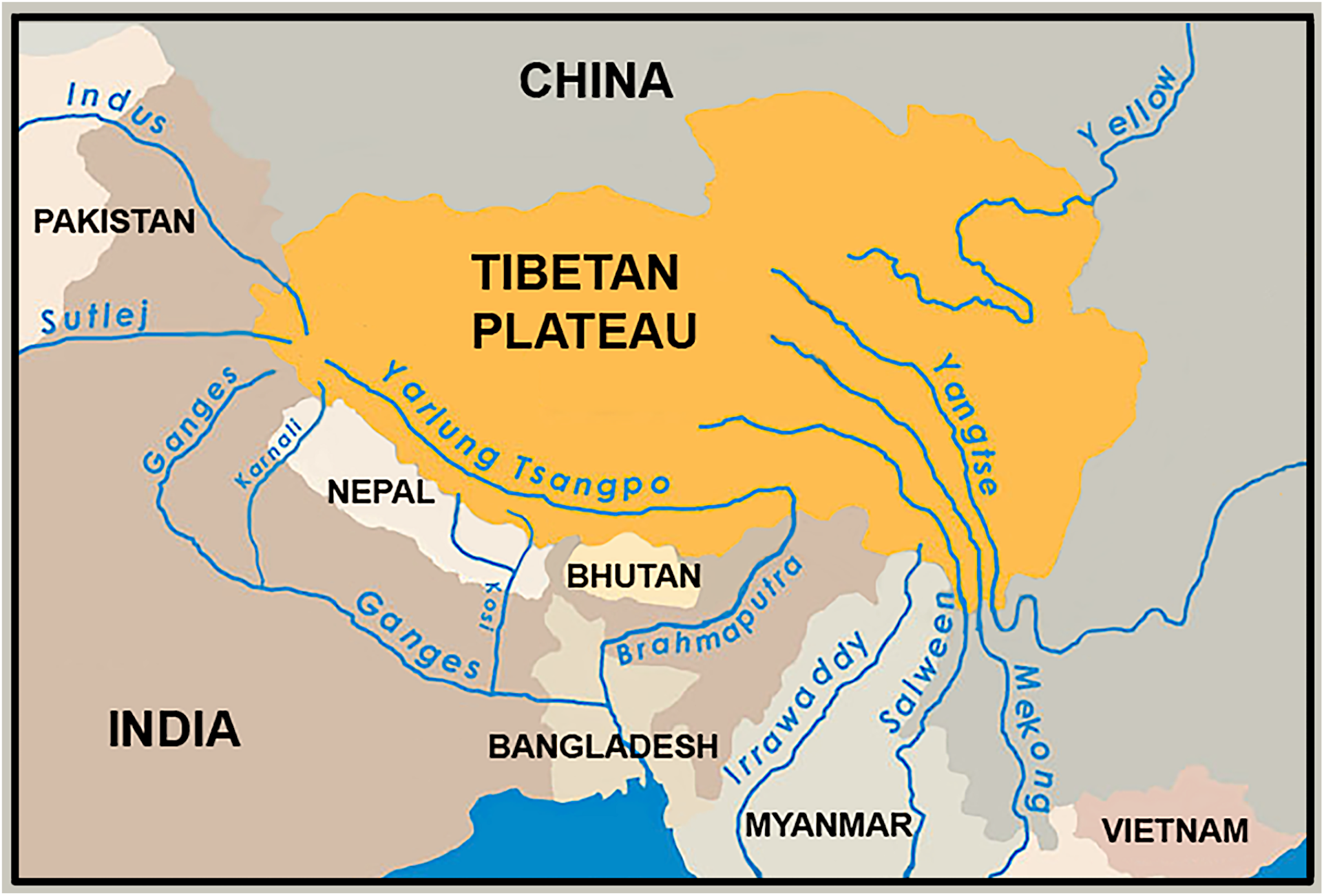
Simplified drainage map of Asia, showing the major basins from which *Schizothorax* were obtained as GENBANK^®^ accessions. Map available via license: CC BY-NC 4.0, and was modified from Fig. 1.17 [86].

The uplift not only established the Himalaya as a topographic entity, but also profoundly influenced the connectivity of its regional rivers [5,6,7]. For example, in late Oligocene–early Miocene (26-19my), the Yarlung-Tsangpo River (YLTR; average elevation 4000m) formed at the southern border of the QTP as a series of lakes with south-flowing tributaries (Fig. 11 in [8]). The lakes eventually coalesced into a westward flowing river (early–mid Miocene; 19–15ma) that eventually reversed flow in response to a continuing but disproportionate uplift. The YLTR was captured on the eastern QTP by the Brahmaputra River in early-mid Miocene (i.e., ~18– 15ma) [7,9], a signature event that seemingly allowed aquatic biodiversity to disperse broadly from east to west in the Himalaya, and to access multiple south-flowing Himalayan drainages.

## Orogeny and biodiversity

The contemporary species richness of terrestrial biodiversity across the QTP clearly reflects the three stratigraphic zones resulting from the uplift. Although this endemism is hypothesized as being driven by tectonic activity, requisite supporting data are infrequent or lacking [10]. The divergence of regional freshwater fishes seemingly occurred when the QTP elevated more than 3,000m in late-Miocene [11], with a potential result being the increased isolation of tributaries. The climatic regime of the QTP was also greatly altered, resulting in strong selection for high-altitude specialists that could seemingly subsist under harsher conditions [12].

Yet, impacts on biodiversity were surprisingly variable. For example, aquatic invertebrates seemingly lose species diversity as elevation increases [13,14], whereas cold-adapted terrestrial species are seemingly buffered by microclimates at higher altitude [15]. However, data for high elevation fishes remain elusive, particularly so in developing countries, an aspect that looms large in the context of monitoring programs that are incomplete and/or non-systematic [16]. This promotes an opportunity for the recognition of cryptic species not as yet identified.

Here, we examined a broadly distributed Asiatic fish (Snowtrout, genus *Schizothorax*), a component of the largest and most diverse of freshwater fish families (Cyprinidae: 1700+ valid species, with 10 recognized subfamilies [17]). Previous research categorized six major biogeographic entities within *Schizothorax* [6]: A Central Asiatic clade, followed in order by five drainage-specific sister-clades – YLTR/ Irrawaddy; Indus; Irrawaddy; Mekong/Salween; Yangtze. However, their phylogenetic interpretation provided an incomplete template for biogeography and evolution of Snowtrout across the entire Himalaya and surrounding regions.

We broadened and extended these results by focusing in particular on two key Himalayan regions previously opaque with regard to freshwater fish diversity: Bhutan (to the Himalayan east) and Nepal (central). Both, in tandem, effectively link the far eastern component of the Himalaya with its western terminus, allowing a more appropriate test of tectonism, orogeny and the evolution of aquatic biodiversity. More specifically, it provides a precise estimate for diversification into eastern, southern, and western Asia, and sets the stage for subsequent climate oscillations in the Quaternary that served to isolate and modify regional biodiversity [18].

## Contemporary significance

The QTP (and surrounding mountains) are termed “the third pole” in that they contain the greatest amount of global snow and ice beyond the Arctic/Antarctic. It is also the headwater source for nine large Asian rivers that collectively provide ecosystem services for >1.5 billion people [19]. Glaciers in the eastern Himalaya, for example, account for an estimated 14.5% of the worldwide total, of which a quarter has been lost since 1970 [20], due largely to a global warming 2x greater than average [2]. Cumulative impacts are recorded as increasing precipitation, decreasing glacial surface area, diminishing snow cover, expanding glacial lakes, and loss of permafrost [21].

The onset of these conditions has (and will continue to) significantly impact the distribution of *Schizothorax*, forcing a migration into higher-elevation streams, as well as concomitant contraction of trailing-edge habitats [22]. These are important considerations given the limitations inherent with *Schizothorax* taxonomy, especially given the strong potential for as-of-yet undiscovered biotic components (e.g. cryptic intraspecific divisions and undescribed species). Regional collaborations that extend across national borders are needed to mitigate climate-change impacts, and to foster cooperative cross-border conservation strategies [21]. Such a regional mandate requires a comprehensive understanding of extant biodiversity, to include inter-specific delineations and the nature by which these juxtapose with both geographical and political boundaries. To do so requires: A) the solidification (and unification) of taxonomies (i.e. to catalogue units for conservation); and B) an established theory for the latter with which to define biogeographic/ phylogeographic expectations. A framework that contrasts these components within a taxonomically and biologically complex genus can be found herein.

## Results

### Sequence data

Our phylogenetic analyses incorporated 139 cytochrome-b (*cyt-b*) haplotypes spanning 1140 bp, with 643 of those monomorphic (=56%), 432 parsimony-informative (38%), and 65 (6%) as singleton polymorphisms. Based on the Bayesian Information Criterion (BIC), the best fitting model of sequence evolution was TN+F+I+G4 (i.e., unequal evolutionary rates among: Purines/pyrimidines, empirical base frequencies, proportion of invariant sites, and a 4-rate category discrete Gamma model). Nepalese and Bhutanese haplotypes (N=53 and N=19, respectively) were compiled across three large, separate drainages within each country.

### Phylogeography of *Schizothorax* across Asia and the Himalaya

To estimate independent groups in our data, we applied a Bayesian approach without *a priori* assumptions (GENELAND V.4.0.3: [23]) and detected evidence for 13 distinct groups. An additional group emerged as a ‘ghost’ lineage but was subsequently disregarded as an artifact of isolation-by-distance [24]. We subsequently explored the manner by which these groups (i.e., N=13 geographic regions; Fig. 2) encapsulated sequence variation by employing an analysis of molecular variance framework (AMOVA; [25]). All were found to be significantly different one from another, with partitions among groups accounting for 78% of the total observed genetic variance (Table 1). Aggregates (= clades) within groups also differed significantly, as did haplotypes within aggregates (Fig. 3). Probabilities for all comparisons were based on 1,000 permutations, with significance gauged by Bonferroni-correction (P<0.005 for 13 groups, p<0.0017 for 29 “aggregates”).

**Figure 2:**
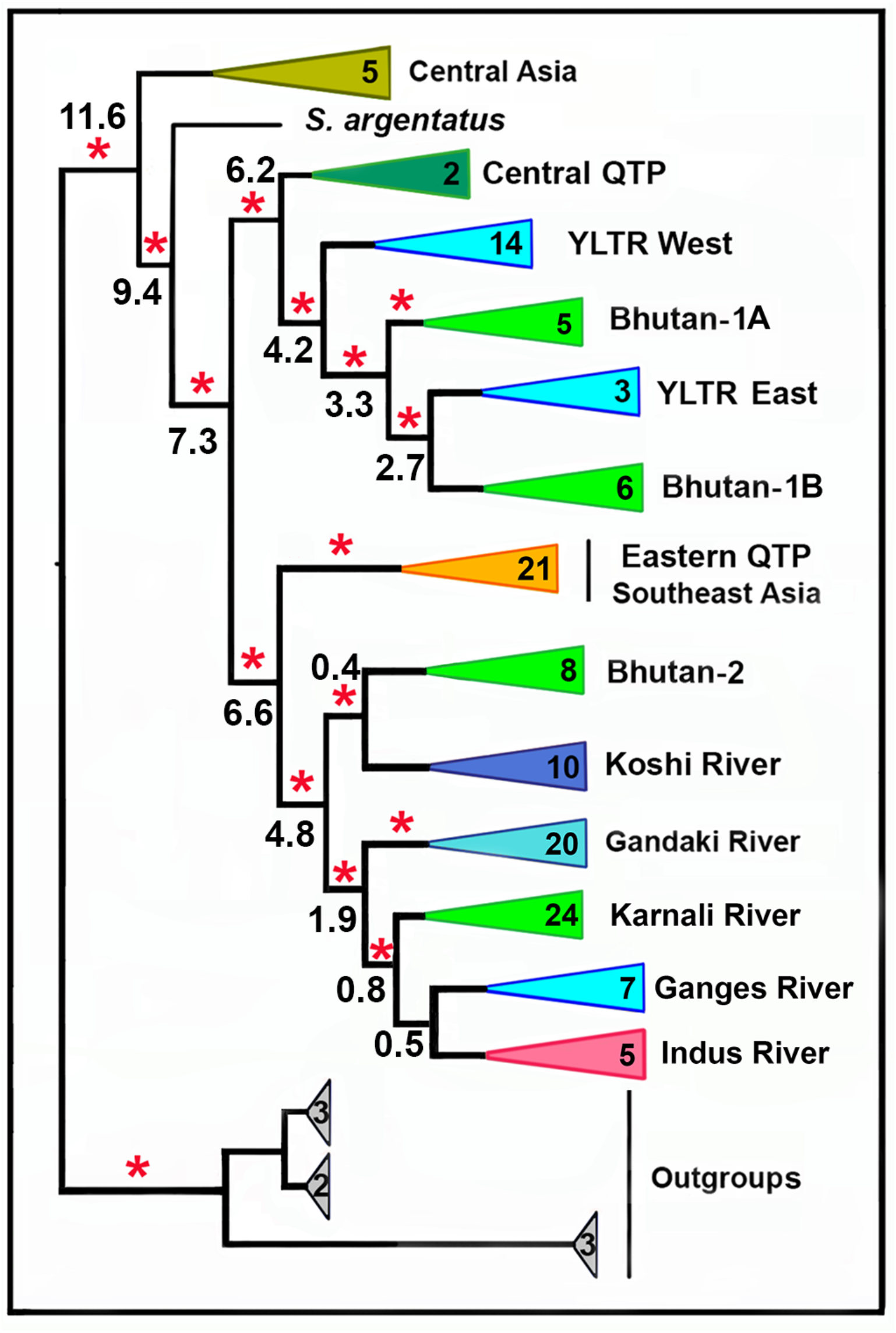
A collapsed maximum likelihood tree reflecting phylogenetic relationships among geographic clades. Data were derived from sequence analysis of the cytochrome-b mitochondrial gene (1140 bp, 139 sequences, redundant haplotypes removed, clades collapsed). Red asterisk = SH-ALRT (Shinodara-Hasegawa approximate likelihood ratio test) ≥ 80% and UFBOOT (ultrafast bootstrap approximations) ≥ 95% (IQ-TREE v.2 Manual: http://www.iqtree.org). Numbers in colored triangles represent species (generalized clades) or individuals (specific clades). Numbers at nodes (N=13) represent RELTIME TIMETREE mean divergence estimates (in Ma; confidence intervals provided in text).

**Figure 3:**
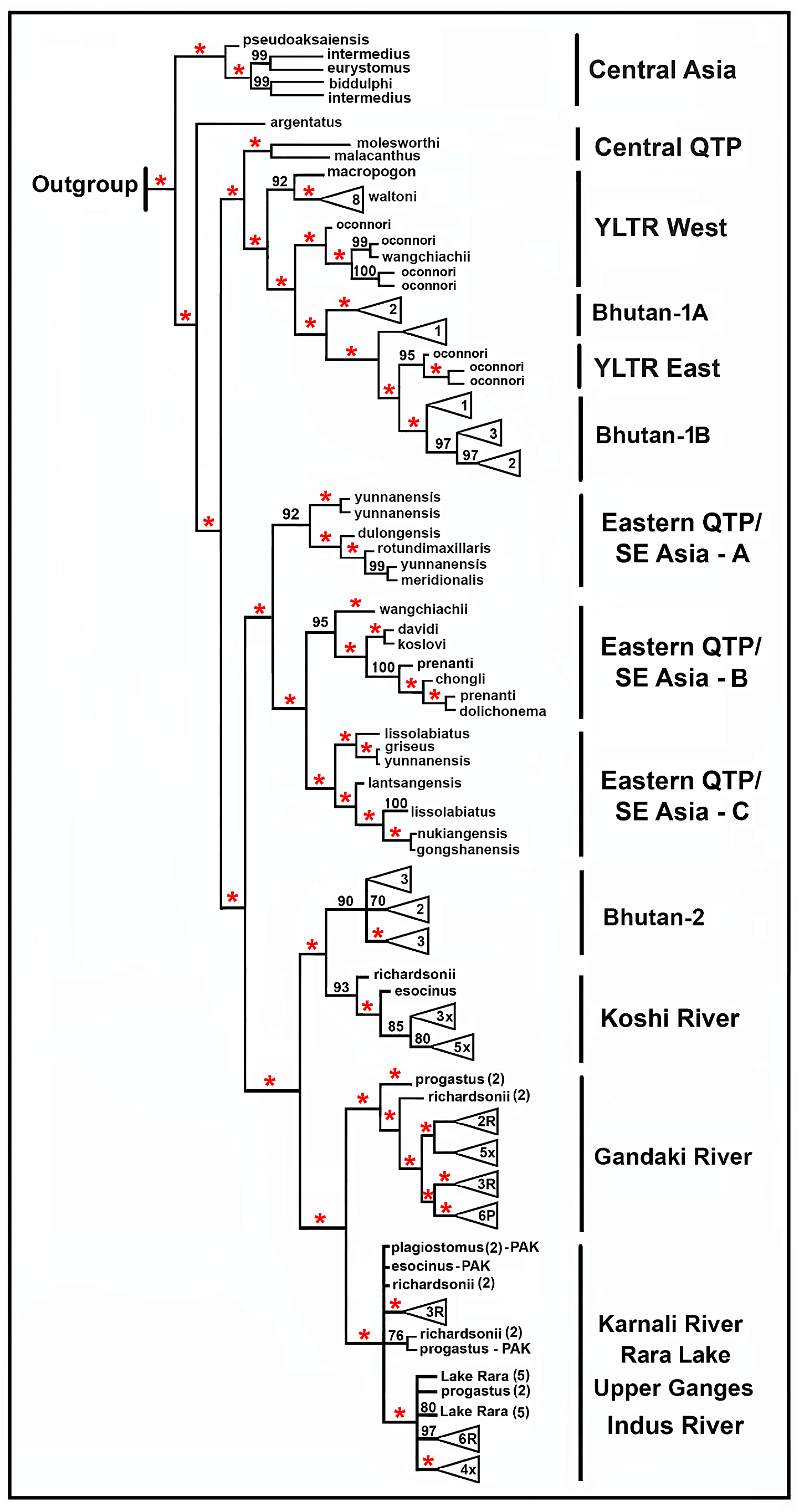
Maximum likelihood tree reflecting phylogenetic relationships among geographic clades. Data were derived from sequence analysis of the cytochrome-b mitochondrial gene (1140 bp, 140 sequences, redundant haplotypes removed). Red asterisk = SH-ALRT (Shinodara-Hasegawa approximate likelihood ratio test) ≥ 80% and UFBOOT (ultrafast bootstrap approximations) ≥ 95% (IQ-TREE v.2 Manual: http://www.iqtree.org). Numbers only at specific nodes represent UFBOOT values. Letters within composite species-triangles at tips of the tree represent species, with R=*S. richardsonii*, P=*S. progastus*, with trailing numbers representing individuals.

**Table 1.** Results of an analysis of molecular variance (AMOVA) based on 130 haplotypes of Snowtrout (Cyprinidae: *Schizothorax*) partitioned into: (1) Geographic regions (=“Among”); Aggregates within clades (=“A/W”); and (3) Haplotypes within aggregates (=“Within”). Source=Grouping; D.F.=Degrees of freedom; S.S.=sums of squares; Var.=variance; %=Percent variance. Probability values for each FST-value (derived via 1000 permutations) are significant at a Bonferroni-adjusted value of p<0.005.

| Source | D.F. | S.S. | Var. | % |
| --- | --- | --- | --- | --- |
| Among | 10 | 3958.7 | 31.4 | 77.9 |
| A/W | 18 | 466.1 | 5.6 | 13.5 |
| Within | 102 | 350.6 | 3.4 | 8.5 |
| Total | 130 | 4775.3 | 40.3 | 99.9 |

We then visualized our groups by deriving maximum likelihood (ML) and Bayesian (BA) trees, each of which resulted in largely consistent topographies (ML tree Figs. 2, 3; Bayesian tree Fig. S1). However, all of our statistical topology tests significantly excluded the BA topology (*p*<0.05), and given this, we thus focus primarily on the ML topology, other than highlighting in a comparative sense the notable differences between the two (Table 2).

**Table 2:** Statistical topology tests comparing maximum likelihood (ML) and Bayesian (BA) phylogenetic results (=Tree). logL=Log Likelihood value; deltaL=logL difference from maximal logL in the set; Bp-RELL=Bootstrap proportions using RELL method (weights sum to 1 across trees); p-KH=*p*-value of one-sided Kishino-Hasegawa test; p-SH=*p*-value of Shimodaira-Hasegawa test; c-ELW=Expected Likelihood Weight (weights sum to 1 across trees); p-AU=*p*-value of Approximately Unbiased test. A plus sign (+) next to a *p*-value denote 95% confidence sets, whereas a minus sign (−) denote a significant exclusion (i.e., the tree is rejected). All tests based on 1000 resamplings using the RELL method.

| Tree | logL | deltaL | bp-RELL | p-KH | p-SH | c-ELW | p-AU |
| --- | --- | --- | --- | --- | --- | --- | --- |
| ML | -9526.83 | 0.0 | 0.986 + | 0.986 + | 1.00 + | 0.978 + | 0.985 + |
| BA | -9582.76 | 55.93 | 0.014 - | 0.013 - | 0.013 - | 0.014 - | 0.015 - |

The majority of nodes in the ML trees [e.g., composite (Fig. 2) and extended (Fig. 3)] were strongly supported on the basis of SH-ALRT (Shinodara-Hasegawa approximate likelihood ratio test) values and ultrafast bootstrap approximations (UFBOOT2). Nodes were likewise supported in the BA tree, with the majority displaying a posterior probability ranging from 0.9-1.0 (Fig. S1).

Two surprising results emerged in both analyses: Bhutanese Snowtrout were consistently non-monophyletic, nesting instead within two temporally and spatially disparate parental clades (i.e., within the Central QTP and the Koshi drainage of Nepal). This indicates multiple, independent colonization of high-altitude habitats within Bhutanese drainages. Nepali Snowtrout, on the other hand, showed strong phylogenetic structuring east-to-west, sequentially diverging in a manner consistent with the broad-scale topographies of riverine networks, and with sub-clades being taxonomically congruent by-and-large. Below, we evaluate results from the ML analysis, but underscore in the discussion those minor discrepancies between ML and BA trees. The specifics of group membership are provided in Table S1.

Groups recovered using genetic clustering approaches (N=13) corresponded well with geographic regions, concordant with groupings apparent in the ML analysis (Fig. 2). These are: (1) Central Asia; (2) Central QTP; (3) YLTR West; (4) Bhutan-1A; (5) YLTR East; (6) Bhutan-1B; (7) Eastern QTP/ Southeast Asia-Clade A (Salween-Irrawaddy drainages); (8) Eastern QTP/ Southeast Asia-Clade B (upper Yangtze drainage); (9) Eastern QTP/ Southeast Asia-Clade C (Mekong-Salween drainages); (10) Bhutan-2 (via Brahmaputra River); (11) Koshi drainage (eastern Nepal); (12) Gandaki drainage (central Nepal); and (13) a composite western Himalaya that included Laka Rara and the Karnali drainage (western Nepal), the upper Ganges (northwest India), and the Indus River Basin (Pakistan). The latter group (i.e., western Himalaya) was additionally partitioned in the BA tree (Fig. S1). For discussion purposes, we subsumed a significantly different single-species clade (*S. argentatus*) within the Central Asia clade (Table S1). A more detailed phylogeny is presented in Fig. 3, with regional composites in Table S1.

Differences between ML and BA-trees are found in the first large geographic cluster that is sister to Central Asia. The clade containing both YLTR regions and Bhutan-1 in the ML-tree was additionally subdivided in the BA-tree: YLTR-West was partitioned by a smaller Bhutan-1A clade, whereas YLTR-East and a larger Bhutan-1B clade were collapsed.

The second large geographic cluster sister to Central Asia in the ML-tree (Fig. 2) has the Eastern QTP/ Central Asia region as a basal node. It has six Himalayan regions as sister groups. The first of these represents eight undescribed Bhutanese samples (=Bhutan-2). The second depicts the Koshi River (eastern Nepal), and both are sister to the Gandaki River group (central Nepal). Of interest is the fact that *S. richardsonii* and *S. progastus* fall together as components of distinct clades, rather than clustering separately by taxonomic designation. A similar result occurs within the Karnali, Ganges, and Indus river drainages. In the BA-tree, the Indus and Ganges drainages appear as distinct clades, whereas they are less so within the ML-tree.

### Temporal divergences

We employed RELTIME to derive major time-calibrated events, 13 of which are depicted in our ML-based tree (Fig. 2; other dates in discussion). The average time for the most basally divergent node (i.e., Central Asia) was 11.6 Ma (CI=15.4-8.7). The two large subclades sister to Central Asia then separated at approximately 7.3 Ma (CI=11.3-4.7). In the first of these, the Central QTP diverged at 6.2 Ma (10.2-3.7), followed by the YLTR West (4.2 Ma; CI=7.4-2.4), then Bhutan-1A (3.3 Ma;CI=6.7-1.5), with YLTR East and Bhutan-1B at 2.7 MA (CI=5.9-1.2).

In the second large subclade sister to Central Asia, the Eastern QTP/ Southeast Asia region diverged at 6.6 Ma (CI=10.7-4.1), followed by Bhutan-2/ Koshi River and Gandaki River/ western Himalayan clades at 4.8 Ma (CI=8.8-2.6). Our Bhutan-2 clade then separated from the Koshi River at 0.4 Ma (CI=0.8-0.2) whereas the Gandaki separated from a composite western Himalayan clade at 1.9 Ma (CI=3.9-0.9) (parsed more thoroughly in Discussion).

## Discussion

### The synergy between tectonism and climate

Orogeny and tectonism drive the evolution of aquatic ecosystems and act synergistically with climate to impact both hydrology [26] and resident freshwater fishes [27]. For example, tectonically derived drainage systems in western North America were gradually shaped over evolutionary time by historic, long-term drought. The result was an erosion of genetic diversity, with endemic fish populations repeatedly collapsing into refugia then expanding outward during onset of more pluvial periods [28]. Episodic flooding, on the other hand, allowed antecedent streams to down-cut, with headwaters subsequently being eroded and basins concomitantly expanding, with large-scale dispersals being promoted [29]. These alterations (i.e., captures, diversions, beheadings; [30]) not only extended the distribution of fishes into adjacent basins [31], but also facilitated hybridization [32,33,34], a process that has confound both taxonomists and managers (and continues to do so).

Flows within basins were often abridged by longitudinal barriers that are vicariant for larger-bodied species yet serve as passive filters for those smaller and more limited in their dispersal [35]. As a result, fish distributions are frequently characterized as a series of biogeographic “islands” within and among basins, each reflecting local expansions, contractions, and extinctions. As such, they provide necessary templates with which to gauge responses by fish to similar tectonic, orogenic, and climatic impositions that have occurred within analogous but less well monitored dendritic ecosystems (as herein).

The Himalaya displays sharp elevational gradients within which there are ample opportunities for diversification. These, in turn, have promoted diversification, that in turn established the region as a global hotspot of biodiversity and endemism [16,36]. Yet, broad phylogeographic studies pertaining to freshwater biodiversity in the Himalaya have been limited, with foci instead on more restricted regions, rivers, and/or mountains [37,38,39,40]. Our perspective was somewhat broader (per [6]), in that we sought molecular evidence for origin and subsequent dispersal of Snowtrout across Asia, as initiated by the continued uplift of the QTP. We were surprised by the robust and coherent biogeographic signal inherent in our data, a consideration that, upon further examination, substantially extends previous results [6].

### Central Asian phylogeography

Our phylogenetic results found five Central Asian species as sister to a significantly different *S. argentatus* (Figs. 2, 3). Of these, S. *pseudoaksaiensis* is distributed in Lake Balkhash and its catchment basin (Kazakhstan, central Asia), as well as the Ili River (northwest China) [41]), as is *S. argentatus*. Similarly, *S. biddulphi* [40] and *S. eurystomus* [42] occur in the Tarim River and tributaries (an endorheic basin in the Central Asiatic Desert of northwest China), whereas the range of *S. intermediu*s is more extensive (i.e., Iran, Uzbekistan, Kazakhstan, Kyrgyzstan). It also occurs in Fig. 3 as two distinct haplotypes.

In the ML tree, the Central Asia clade diverged at 11.6 Ma (CI=14.3-6.2), an estimate comparable with previous results (11.4-10.5 Ma; [6]). Our timetree estimation for S. *argentatus* (9.4 Ma; CI=14.3-6.2) also corroborates earlier estimates (9.6-8.9 Ma; [6]). It thus seems reasonable to recognize Central Asia as the center of origin for schizothoracins, although this node was not placed as such in the statistically less supported BA-tree (Table 2; Figure S1).

The next major split in our ML phylogeny defines two large nodes. The first encompasses components of the QTP (save those in the far east) plus Bhutan (immediately south), whereas the second incorporates the eastern QTP/ Southeast Asia plus those Himalayan components that dispersed south via the Brahmaputra River then west. Our timeline is 7.3 Ma (CI=11.3-4.7), again analogous with previously estimated values (8.3-7.6 Ma; [6]). We discuss both large clades and their components below.

### Phylogeography of the Central QTP, YLTR, and “old Bhutan”

The two species in the Central QTP (*S. molesworthi and S. malacanthus*) separated in early Miocene (6.2 Ma; CI=10.2-3.7), whereas the YLTR West diverged late Pliocene at 4.2 Ma (CI=7.3-2.4). Here, *S. macropogon* (3.2 Ma; CI=6.6-1.5, late Pliocene) is sister to *S. waltoni* which subsequently differentiated into western and eastern groups that are detectable via microsatellite (msat) DNA but not *cyt-b* [43]. It is, in turn, sister to a second YLTR West clade (*S. oconnori* + *S. wangchiachii*).

The evolutionary history of *S. oconnori*, a species recognized as endemic to the YLTR [41], underscores the role played by the QTP uplift in initiating late Miocene divergence. During this process, *S. oconnori* was subsequently separated into a western component above a precipitous canyon in the YLTR, and an eastern component below [44]. Indeed, this hypothesis is clearly sustained in our composite analyses (SH-ALRT, UFBOOT, AMOVA), with the western clade (to include *S. wangchiachii*) not only significantly different from eastern *S. oconnori* (YLTR-East), but also separated from it by an undescribed clade from Bhutan (i.e., Bhutan-1A; Fig. 3). A second Bhutanese clade (Bhutan-1B) is sister to YLTR-East.

This seemingly indicates two distinct dispersals by *S. oconnori* from the ancestral YLTR. One involved Bhutan-1A (the western clade), whereas a second represents the eastern clade (Bhutan-1B). Both Bhutanese clades are currently unverified as to species but can seemingly be allocated to *S. oconnori*. Quaternary climatic oscillations on the southern YTP not only yielded evolutionarily significant units (ESUs) in both *S. waltoni* and *S. oconnori*, but also in Bhutanese *Schizothorax* as well. Again, the BA-tree differs somewhat in that the YLTR lineages are further subdivided. The YLTR-West region is broken by the Bhutan-1A clade, whereas Bhutan-1B (the larger clade) becomes collapsed within the YLTR-East region.

### Phylogeography of the Eastern QTP and Southeast Asia

The Eastern QTP/ Southeast Asia group is represented by three significantly different clades (Fig. 3). The deepest divergence (i.e. Clade A from the remainder; Table S1) encompasses six species from the Irrawaddy/ Salween rivers, with a timetree estimate at 5.6 Ma (CI=9.7-3.0). The second (Clade B; Table S1) consists of seven species from the Yangtze River (at 1.7 Ma; CI=3.5-0.8), whereas the third (Clade C; Table S1) involves eight species from the Mekong/ Salween drainages. Divergence in the Mekong was seemingly driven by late Pliocene orogenesis [37], an estimate that is congruent with our derived timeline (at 2.4 Ma; CI=5.0-1.2). Three species within the upper Salween River (i.e., *S. gongshanensis, S. lissolabiatus, S. nukiangensis*) display low genetic differentiation and frequent haplotype sharing as a by-product of Quaternary glaciations [39]. However, several species in clade B (such as S*. davidi, S. koslovi, S. chongli, S. gongshanensis*) have yet to be fully described or documented, other than in general terms [40].

### Dispersal of *Schizothorax* into Bhutan, Nepal, and western Himalaya

The YLTR originates on the QTP, flows eastward then abruptly south as it passes through the Namche Barwa Syntaxis [5] where it was subsequently captured by the Brahmaputra River [7]. As such, it represents an important dispersal route from the QTP for freshwater species, and also an important phylogenetic bookmark in that it provides an evolutionary transition between species on the QTP versus those that diverged more recently, subsequent to westward dispersal via the Lower Brahmaputra followed by northward movements into Himalayan drainages that have previously down-cut to the south.

Our samples from Nepal form sister clades, each representing a distinct drainage, all of which differ significantly amongst themselves, and from Southeast Asian clades (Fig. 2). Here, additional samples from Bhutan (Bhutan-2; N=8) form a distinct clade sister to the eastern-most drainage of Nepal (Koshi River Basin clade; N=11, Fig. 4, Table S1), thus effectively underscoring the east-to-west transition mediated by dispersal from the lower Brahmaputra. Samples from Bhutan-2 are currently not identified as to species, whereas those in the Koshi clade are referred to *S. richardsonii, S. esocinus*, and *S. progastus.* The morphological similarities between *S. richardsonii* and *S. progastus* are such that their discrimination in the field is problematic [45], particularly at early life history stages. One hypothesis is that these inconsistencies may contribute to the apparent phylogenetic assimilation of both species within several Koshi, Karnali, and Gandaki river clades (Fig. 3).

**Figure 4:**
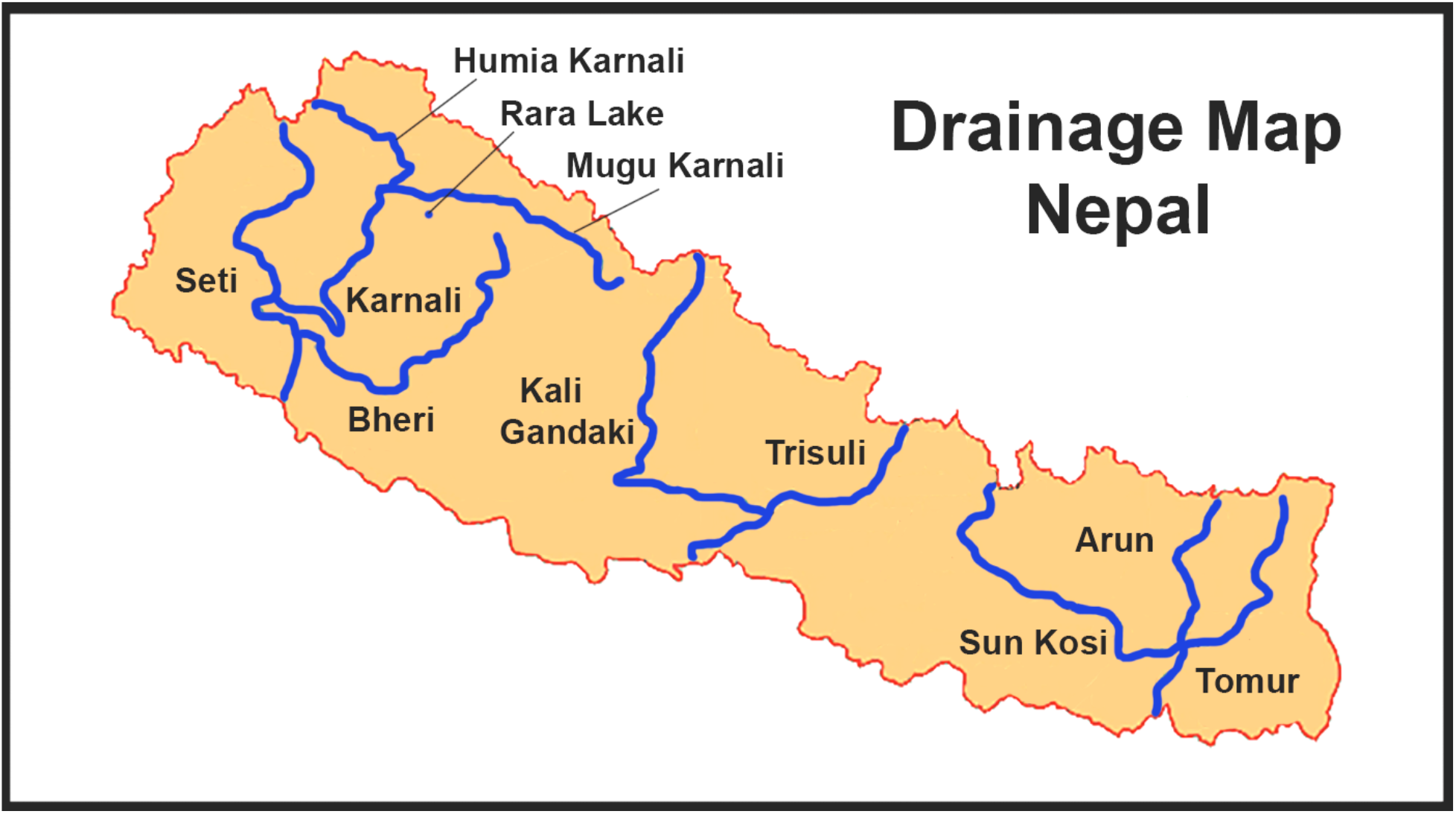
Drainage patterns of Nepal (Central Himalaya). Three major river basins merge with the Ganges River in the lowlands of India. Lake Rara (northeast Nepal) drains into the Karnali River. Details on *Schizothorax* localities are provided in Appendix 1.

Samples from the Gandaki and the Karnali rivers each form monophyletic clades as well. The Gandaki (N=20) is partitioned among *S. richardsonii* (three clades), S. *progastus* (two clades), and one representing a mixture of the two (Fig. 3). Samples from the Karnali River Basin (far western Nepal, N=24; Fig. 4) fall within a composite that encompasses the Upper Ganges (N=7) and Indus rivers (N=5; western Himalaya). Species from the Gandaki and Indus river basins are *S. plagiostomus* and *S. progastus*, with *S. esocinus* also from the Indus. Once the Koshi, Gandaki, and Karnali rivers reach the lowlands of India, they drain into the Ganges which continues eastward to the Brahmaputra River then to the Bay of Bengal. (Fig. 1). The Indus, however, now drains westward to the Arabian Sea.

Additional discrepancies appear between ML and BA-trees with regard to discrimination among western Himalaya regions. Specifically, the BA-tree separates the Indus and Ganges into separate clades, as sister to (and differentiated) from the Karnali River/ Rara Lake clade.

### Phylogeography of Rara Lake, Nepal

Three morphologically differentiated and reproductively isolated *Schizothorax* (i.e., *S. macrophthalmus, S. nepalensis, S. raraensis*) occupy separate trophic niches in Lake Rara (Karnali River Basin, northwestern Nepal: [46]), with *S. raraensis* and *S. nepalensis* listed as critically endangered (IUCN, 2017; http://www.iucnredlist.org/details/168564/0/). Of the three, only *S. nepalensis* could be differentiated using mtDNA and, given this, it was hypothesized [47] that *S. macrophthalmus* and *S. raraensis* shared mtDNA haplotypes via hybridization. Similarly, two morphologically distinct Snowtrout in the upper Karnali River (i.e., *S. richardsonii* and *S. progastus*) could not be discriminated as well, with each similarly hypothesized as exemplifying interspecific mitochondrial exchange [47]. Our mitochondrial marker could not resolve this hypothesis, thus necessitating further nuclear analysis so as to unequivocally test the role of hybridization as a source of phylogenetic discordance. Such an examination, however, is further complicated by polyploidy and ploidy variance within *Schizothorax.*

The geologic age of Lake Rara is estimated as ~60kya [48]. Its endemic species do not form a monophyletic clade, but cluster instead with tributaries of the Karnali River that drain the lake (Fig. 3). Haplotypes of *S. richardsonii* and *S. progastus* in the Karnali River Basin did not differ significantly from one another yet were significantly different when compared with conspecifics in the Koshi and Gandaki rivers [47]. Our analyses confirmed these results (Fig. 3). Speciation in the Karnali River Basin could thus be poorly resolved with respect to mitochondrial divergence, with a genetic signal likely obscured by the limited framework within which ancestral alleles could sort, allowing novel haplotypes to arise. More refined molecular approaches, such as reduced-representation genomics [32,33,34,49,50], could potentially provide the necessary resolution, although it will again be compounded by the rampant polyploidy found in the subfamily Cyprininae.

### Dispersal of Snowtrout into western Himalayan tributaries

Recent studies [6,37,51,52] suggest the current distribution of Snowtrout has been shaped by late Miocene uplift of the QTP, with vicariant events occurring within headwaters of several rivers in the Eastern QTP/ Southeast Asia clade. However, these are localized perspectives rather than broadly regional, due largely to conflicting interpretations regarding the evolution of Himalayan paleodrainages, particularly those in the western Himalaya. A second confounding factor is a deficit of molecular samples from the Central Himalaya (a crucial link between eastern and western components of the Himalaya).

The southern QTP and western Himalaya are now drained by the contemporary Indus River which extends south and west from the Karakorum Range, rather than from the High Himalaya. Given its western placement, the geomorphic history of the Indus is a key component with regards to the biogeography of *Schizothorax* in the Himalaya. The relationship of the Indus with the Ganges immediately to the east is unsettled, with several proposed hypotheses [52].

Relative stasis between the two is the most conservative, whereas others involve various stream-capture scenarios, such as: The Ganges initially flowing westward to the Indus then to the Arabian Sea; The Indus flowing eastward into the Ganges then to the Bay of Bengal; and lastly, an earlier capture by the Ganges of four western Himalaya rivers (i.e., Jhelum, Chenab, Ravi, Sutlej) that are now incorporated within the contemporary Indus Basin.

The latter hypothesis is supported by isotopic and seismic data extracted from the detrital fan of the Indus River, as deposited for >30 Ma in the Arabian Sea. These data document a major shift at ~5 Ma, with earlier Himalayan deposition being superseded by those from the Indus suture zone to the north [53]. This shift documents the capture by the Indus of several large Punjabi rivers (i.e., Jhelum, Chenab, Ravi, Sutlej), as represented by the current topography of the basin. However, it is not without controversy. A subsequent comparison of sediments from northern Pakistan with those of the Indus Fan suggests the documented transition was a response to differential erosion rates in the source region (the Indus) rather than drainage rearrangements, as above [52].

The strong east-west divergence among Himalayan clades found in our data represents biological data from which to interpret the geomorphic evolution of Himalayan drainage systems. In this sense, our phylogeographic results are a biological corroboration of the capture by the Indus of those drainages previously allocated to the Ganges. *Schizothorax* from the Lower Ganges River diverged from a clade representing the Upper Ganges Basin and the Indus Basin. Similar patterns have been noted for Indus and Gangetic dolphin [54]. These data also suggest *Schizothorax* in the western Himalaya (i.e., Koshi, Gandaki, and Karnali) was isolated prior to the capture of the Jhelum by the Indus.

### A confused taxonomy for *Schizothorax*

Several species were found to be polyphyletic in our analyses, with individuals forming biogeographically defined clusters that differed significantly from one another. In addition, multiple cases were found where samples within the same region also failed to cluster with conspecifics (e.g., Central Asia; Eastern QTP/Southeast Asia; Koshi; Gandaki; Karnali; Ganges).

This phenomenon highlights several trenchant issues manifested within the schizothoracines: Incongruence between molecular and morphological phylogenies [6,37,51], and convergent evolution (e.g., juxtaposition of trophic and spatial niches across species [55]). An observed lack of association between morphological and molecular characters may be due either to incomplete lineage sorting, or rapid diversification [18]. If the latter, then mtDNA as a molecular marker will be incapable of resolving these issue [6]. These issues are amplified, and biological interpretations consequently hindered, by the potential for DNA sequences within online databases to be incorrectly annotated with regards to taxonomy [56].

## Conclusions

*Schizothorax* in the Himalaya is a monophyletic clade that originated in Central Asia, subsequently dispersed southward into Bhutan via down-cutting rivers, then westward across the Eastern QTP and into Southeast Asia. Subsequent capture of the YLTR by the Brahmaputra allowed for an east-to-west dispersal below (i.e., south of) the Himalaya, with subsequent movements northward into sequential Himalaya drainages, first Bhutan, then Nepal, followed by the western Himalaya. Our molecular data also serve to corroborate western drainages previously affiliated with the Ganges as being subsequently captured by the Indus River.

Many morphologically recognized species have seemingly diverged substantially across basins, suggesting the potential for cryptic speciation and/or undescribed relictual biodiversity. Additionally, more detailed molecular studies would promote conservation planning at the regional scale but will also require the analysis of populations now geographically isolated across the Himalaya. This, in turn, will necessitate cooperation/ collaboration among neighboring countries, particularly given the recognized (as well as predicted) impacts of climate change on the Third Pole in general, and *Schizothhorax* in particular [22].

## Methods

### Ethics Statement

All methods were performed in accordance with relevant guidelines and regulations. Bhutanese Collecting permits were in conjunction with the National Research & Development Centre For Riverine and Lake Fisheries (NRDCR&LF), Ministry of Agriculture & Forests (MoAF), Royal Government of Bhutan. The export of fin clips was authorized through a Material Transfer Agreement (MTA) provided by the Bhutan Agricultural and Food Regulatory Authority (BATRA). Sampling protocols were approved by the University of Arkansas Institutional Animal Care and Use Committee (UA_IACUC_17064).

### Drainages of Nepal and Bhutan

Three major drainage basins in Nepal (east-to-west: Koshi, Gandaki, and Karnali rivers; Fig. 4), support eight tributaries, several of which originate on the QTP, although we found no evidence for historic dispersal of *Schizothorax* directly into Nepal. Mountainous slopes in the Koshi River are more eroded and abruptly elevated, given that the monsoon has a stronger effect on eastern Nepal than further west. Central Nepal is drained by the Gandaki Basin, and its western tributary (Kali Gandaki) is recognized as a vicariant east-to-west break. Western Nepal, on the other hand, is drained by the Karnali Basin, and includes one of our study sites (i.e., Lake Rara, a RAMSAR site with a surface area of 10.6 km^2^ at 2,990m elevation). All three basins drain south into the east-flowing Ganges River that subsequently joins with the Brahmaputra River and terminates in the Bay of Bengal.

Bhutan, on the other hand, displays five major north-south river systems (Fig. 5). The Amo Chhu, Wang Chhu, and Punatsang Chhu (=Sankosh) drain western Bhutan, with the Amo Chhu and Wang Chhu descending from the QTP and flowing southeasterly through west-central Bhutan. The Punatsang Chhu drains the Great Himalayan Range. All three subsequently pass into West Bengal (India).

**Figure 5:**
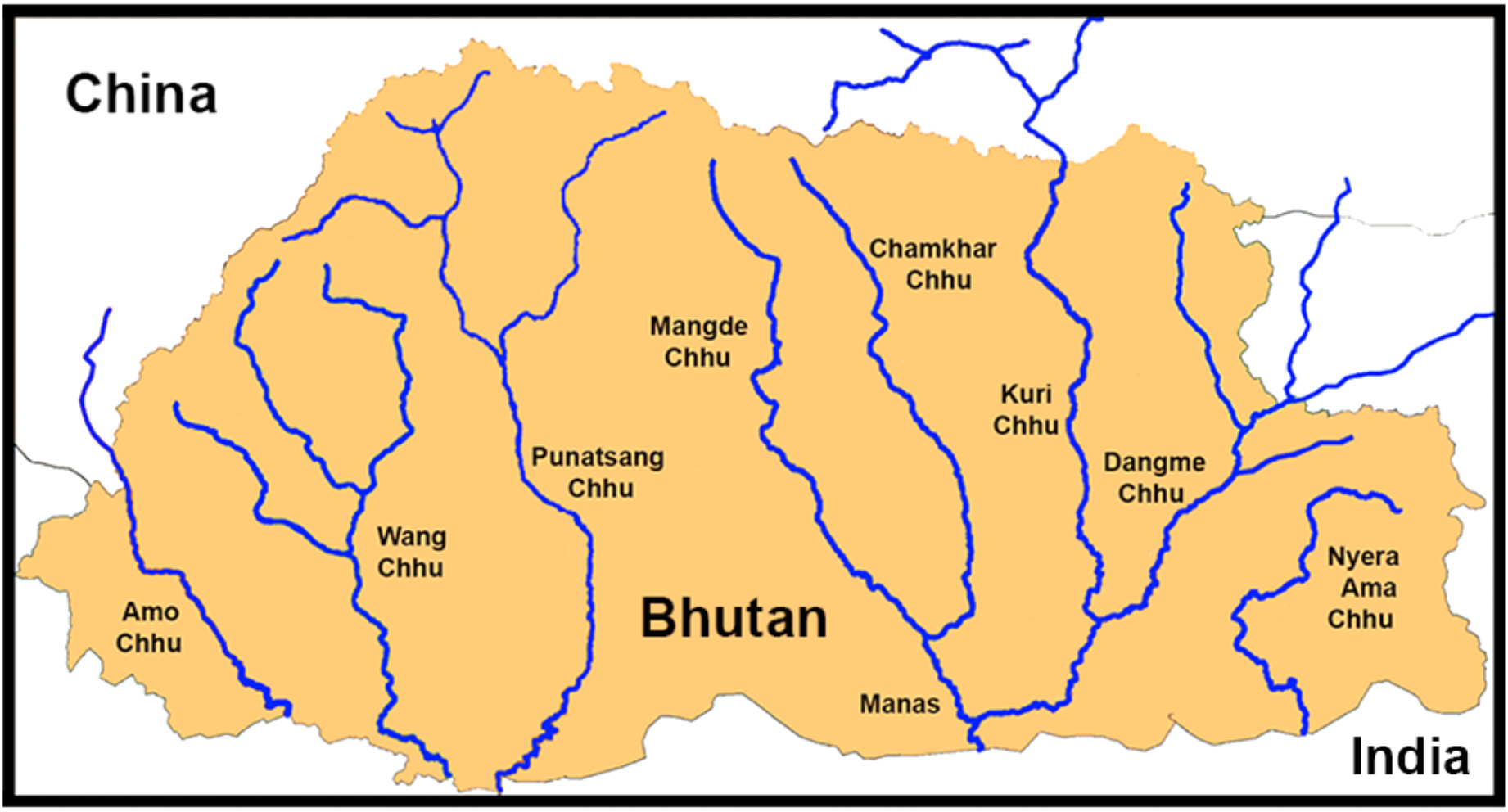
Drainage patterns of Bhutan (Central Himalaya). Six major rivers drain southward into the Brahmaputra River (northwest India). Samples were obtained from: Amo Chhu, Wang Chhu, Punatsang Chhu/Sunkosh, and Mangde Chhu/Manas. Details on *Schizothorax* localities are provided in Appendix 1.

The Manas and its several major tributaries drain central-eastern Bhutan. There, the Dangme Chhu flows southwesterly from Arunachal Pradesh (India), whereas the Kuri Chhu (from the QTP), the Chamkhar Chhu, and the Mangde Chhu drain central Bhutan. All Bhutanese basins drain eventually into the Brahmaputra River (India), which curves south and west through Bangladesh to merge with the Ganges River, then to the Bay of Bengal (Fig. 1).

### The origin of *Schizothorax* and its outgroups

Snowtrout (Fig. 6) is an ideal organism to decipher the geologic processes that drove historic and contemporary fish distributions in the Himalaya. It contains a diverse array of species distributed across deserts, lakes, and rivers of the QTP, as well as the snow-fed tributaries of the Himalaya [57,58]. Some 28 *Schizothorax* species are recorded in the Himalayan, sub-Himalayan, and Tibetan regions [59].

**Figure 6:**
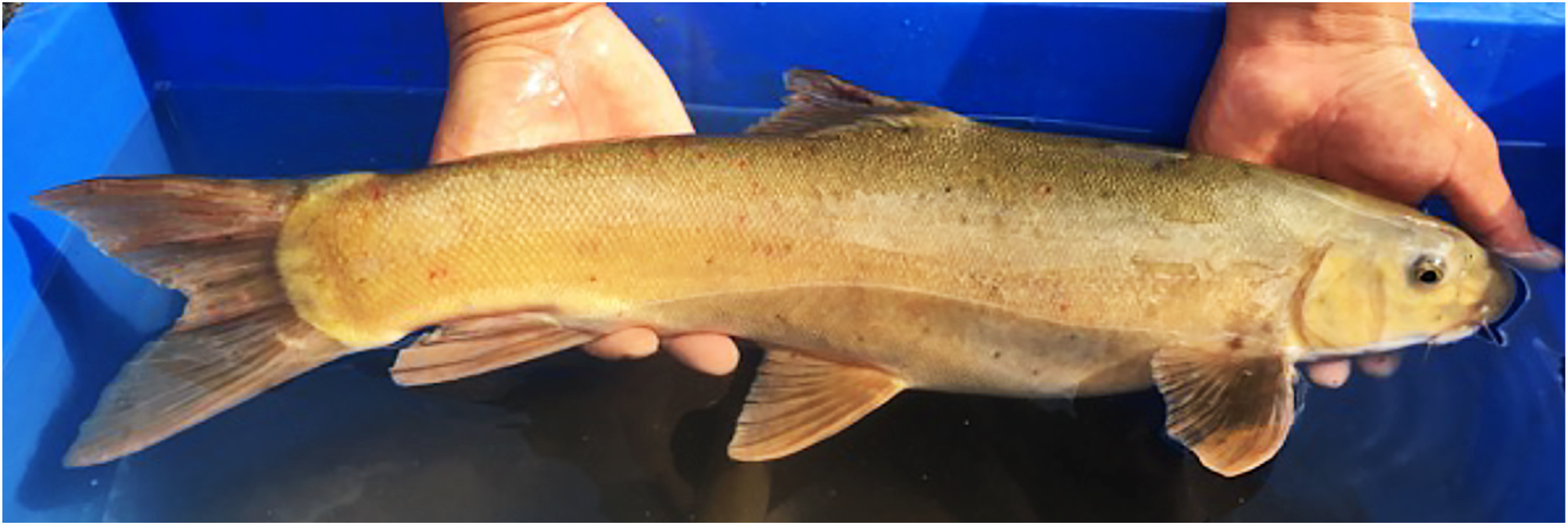
*Shizothorax sp.* (potentially *S. richardsonii*) captured at the confluence of the Berti Chhu and Mangde Chuu, Bhutan.

Several hypotheses have been offered to explain the origin and dispersal of *Schizothorax* across the Himalaya and Asia. Previous researchers [60] suggested the subfamily Schizothoracinae initially dispersed into the QTP from southeastern Asia, and subsequently became isolated by orogeny and climate change. Others [61] proposed that tectonism isolated a primitive barbin clade on the Tibetan Plateau which then served as progenitor for a contemporary *Schizothorax* radiation.

A recent revision of the subfamily Cyprininae [62] was based on a combined series of molecular markers: *cyt-b* (N=791), mitogenomes (N=85), and nuclear RAG-1 sequences (N=97)]. Results retained *Schizothorax* within the monophyletic Tribe Schizothoracini that (depending upon the marker) evolved prior to or after the Barbini. Thus, in selecting outgroups, we avoided Barbini and instead fixed on tribes identified as evolving prior to the Schizothoracini. These included (in sequence): Torini (i.e., *Tor* and *Neolissochilus*), Smilogastrini (i.e., *Puntius, Pethia*), and Acrossocheilini (i.e., *Acrossocheilus*, the most immediate outgroup) (Table S1).

### Sampling and the amplification/ sequencing of mtDNA

Nepali samples (N=53) represented five *Schizothorax* species. Three [i.e., *S. macrophthalmus* (N=3), *S. nepalensis* (N=4), and *S. raraensis* (N=3); total N=10] represented a landlocked species-flock within Rara Lake (Karnali River drainage; Fig. 4). Two other species [*S. richardsonii* (N=28), and *S. progastus* (N=15)] are riverine and were sampled from (west-to-east): Karnali River (N=12/N=2, respectively), Gandaki River (N=11/9), and Koshi River (N=5/4).

Nepali samples were supplemented with those from Bhutan (N=19) collected from four drainage basins (west-to-east): Wang Chhu (N=1), Punatsang Chhu (N=13), Mangde Chhu (N=3), Dangme Chhu (N=1), Nyera Ama Chhu (N=1); Fig. 5). Genomic DNA was extracted using the Qiagen DNeasy kit, per manufacturer’s instructions. In addition, we accessed 67 mtDNA *cyt-b* sequences representing 36 species (28 *Schizothorax* plus eight outgroups [*Tor* (N=2), *Neolissochilus (*N=1), *Puntius* (N=1), *Pethia* (N=2), *Acrossocheilus* (N=2); Table S1] (National Institute of Health GENBANK^®^, https://www.ncbi.nlm.nih.gov/genbank/). Total number of individuals = 139.

### Polyploidy

The subfamily Cyprininae displays rampant polyploidy (>400 species; [62,63], an important consideration in molecular studies, in that methodological accommodations across domains (e.g., population genetics) are lacking. This is in particular problematic within a phylogenetic context, given the greater redundancy of nuclear genes in polyploid genomes. All may potentially diverge one from another over evolutionary time, such that results could differ markedly depending upon which ‘sub-genome’ is accessed when polyploid molecular data are phylogenetically evaluated [64].

Given this, phylogenetic analyses of polyploid species are often limited in practice either to mtDNA data (due to its haploid nature), or nuclear genes cloned into bacteria and subsequently isolated as single-copy [such as Recombination Activating Gene 1 (RAG-1); [62], though the latter is recognized as more conservative evolutionarily, and without a timetree as an option [65]. With regards to *Schizothorax,* polyploidization may represent an evolutionary mechanism that facilitated an invasion of high-elevation habitats [63].

### Phylogenetic analyses

We amplified mtDNA *cyt-b* using published primers [66], with products being enzymatically purified, sequenced using BIGDYE (ver.3.1) chemistry (Applied Biosystems Inc. [ABI], Forest City, CA), and analyzed on an ABI Prism 3700 Genetic Analyzer (W.M. Keck Center for Comparative and Functional Genomics, University of Illinois, Urbana/Champaign). Sequences were edited/aligned using SEQUENCHER v.5.3 (Gene Codes Inc., Ann Arbor, MI), with nucleotide composition and uncorrected p-distances derived among groups (MEGA X; [67].

We then developed a dataset that included only variable sites (N=365bp: DNAsp v.6; [68]) which was subsequently employed in a Bayesian approach (GENELAND V.4.0.3: [23,69]) to infer the number of distinct groups (*K*) and designate group membership. Ten runs of 100,000 MCMC iterations and a thinning interval of 100 were used, with *K* varying between 1 and 15, but without use of optional geo-referencing, due to the absence of coordinates for most sequences in the GENBANK^®^ metadata. The estimated number of populations (K) was defined using the maximum posterior estimate. We also conducted an analysis of molecular variation (AMOVA: ARLEQUIN v.3.5.1.2; [25]) so as to evaluate geographic “groups” (as informed by GENELAND and represented in Fig. 2). Here, we defined “populations” (= distinct clades) as being haplotypes within groups [70]. Probabilities were ascertained using 1,000 permutations, and statistical significance was assigned using a Bonferroni-correction.

We derived phylogenetic clades using a maximum likelihood (ML) analysis (IQ-TREE v2; [71]), with the best-fitting substitution model generated via MODELFINDER, with flexible rate heterogeneity (e.g. gamma-distributed) across sites [72]. This was accomplished by exploring model space hierarchically, adding parameters (e.g. rate categories) sequentially until the model failed to improve. Support for the resulting tree, inferred using the selected model, was explored using 1,000,000 ultrafast bootstrap approximations (UFBOOT2; [73]). We also assessed branch support using the SH-ALRT test (Shinodara-Hasegawa approximate Likelihood Ratio Test), with values ≥ 80% indicating strongly supported clades, whereas those for UFBOOT are ≥ 95% (IQ-TREE v.2 Manual: http://www.iqtree.org).

To employ a Bayesian approach, we first derived an XML input file using the graphical user interface (i.e., BEAUTY) for the phylogenetic software suite BEAST (v2.61; [74]), using a relaxed molecular clock with an uncorrelated lognormal rate distribution and a Yule process prior. Each analysis was run for 100 million generations, with samples taken every 1,000^th^ iteration, following a burn-in of 20%. Effective sample size (ESS) was verified as >200 across all parameters with MCMC chain convergence visually inspected using TRACER [75]. Trees from three independent runs were combined (via LOGCOMBINER) and represented as a maximum clade credibility (MCC in TREEANNOTATOR: [74]), with nodal support representing posterior probabilities. The resulting tree was visualized using FIGTREE (v.1.4.3; [76]).

Topologies from ML and BA analyses were statistically compared by assessing each with respect to the original sequence alignment using IQ-TREE2 v2 [70]. To do so, we examined support for each using RELL approximation with 10,000 replicates using: (a) Raw change in log-likelihoods; (b) Bootstrap proportions [77]; (c) Kishino-Hasegawa test [78]; (d) Shimodaira-Hasegawa test [79]; (e) Approximately Unbiased test [80]; and (f) Expected Likelihood weights [81].

### Developing a timetree

For molecular dating, we employed a rapid, high-performance ML-based model (RELTIME, MEGA X; [67]) that incorporates a relative rate framework (RRF) within which the evolutionary rates of sister lineages are contrasted against a minimum rate change, as diagnosed between evolutionary lineages and their respective descendants. The computational speed and accuracy of the model is appropriate for the analysis of large data sets (>100 ingroup sequences; [82]). The procedural steps for RELTIME are described in [83]. The evolutionary model for our ML-based phylogenetic tree (i.e., TN+G4+I) was also employed in RELTIME.

We estimated a divergence time based upon the earliest fossil recognition for the Schizothoracini (e.g., Oligocene: *Paleoschizothorax qaidamensis* [84]). The stratigraphic formation within which this Qaidam Basin fossil occurred was gauged as Eocene-to-Oligocene [85], and given this, we chose as our divergence time the Oligocene initiation (at 33Ma).

## Acknowledgments

Nepali samples were obtained from the University of Kansas Natural History Museum (http://www.kansastravel.org/lawrence/kansasnaturalhistory.htm) and represent a subset of those collected by DRE during completion of a 1996 Fulbright Award. We thank Curator Leo Smith and Collection Manager Andrew Bentley for their prompt response and support. We also acknowledge the National Research Centre for Riverine and Lake Fisheries (NRCRLF, Bhutan) for providing logistic support and personnel with which to sample fishes. Analytical resources were provided by the Arkansas Economic Development Commission (Arkansas Settlement Proceeds Act of 2000) and the Arkansas High Performance Computing Center (AHPCC). This research was made possible through generous endowments to the University of Arkansas: The Bruker Professorship in Life Sciences (MRD), the 21^st^ Century Chair in Global Change Biology (MED), Distinguished Doctoral Fellowship awards (TKC, ZDZ), and Graduate Teaching Assistantships (KW, TKC, ZDZ), all of which provided salaries and/or research funds to complete this study. Components of this study also represented a research project (ZA) completed at the University of Arkansas Honors College.

## Author Contributions

BR, MRD, MED, DRE, KW, and ST conceived and designed the study; DRE, KW, CL, GPK, PN, SN, SD, TKC, and ZDZ collected samples in the field; BL, ZA and MED analyzed the data; BL, MRD, MED, and ZA wrote the article; All authors discussed the results and commented on the manuscript.

## Additional Information

### Competing Interests

The authors declare no competing interests.

## Data Availability Statement

mtDNA data: Dryad Digital Repository (http://dx.doi.org/upon-acceptance).

---

**Supplementary Table S1:**
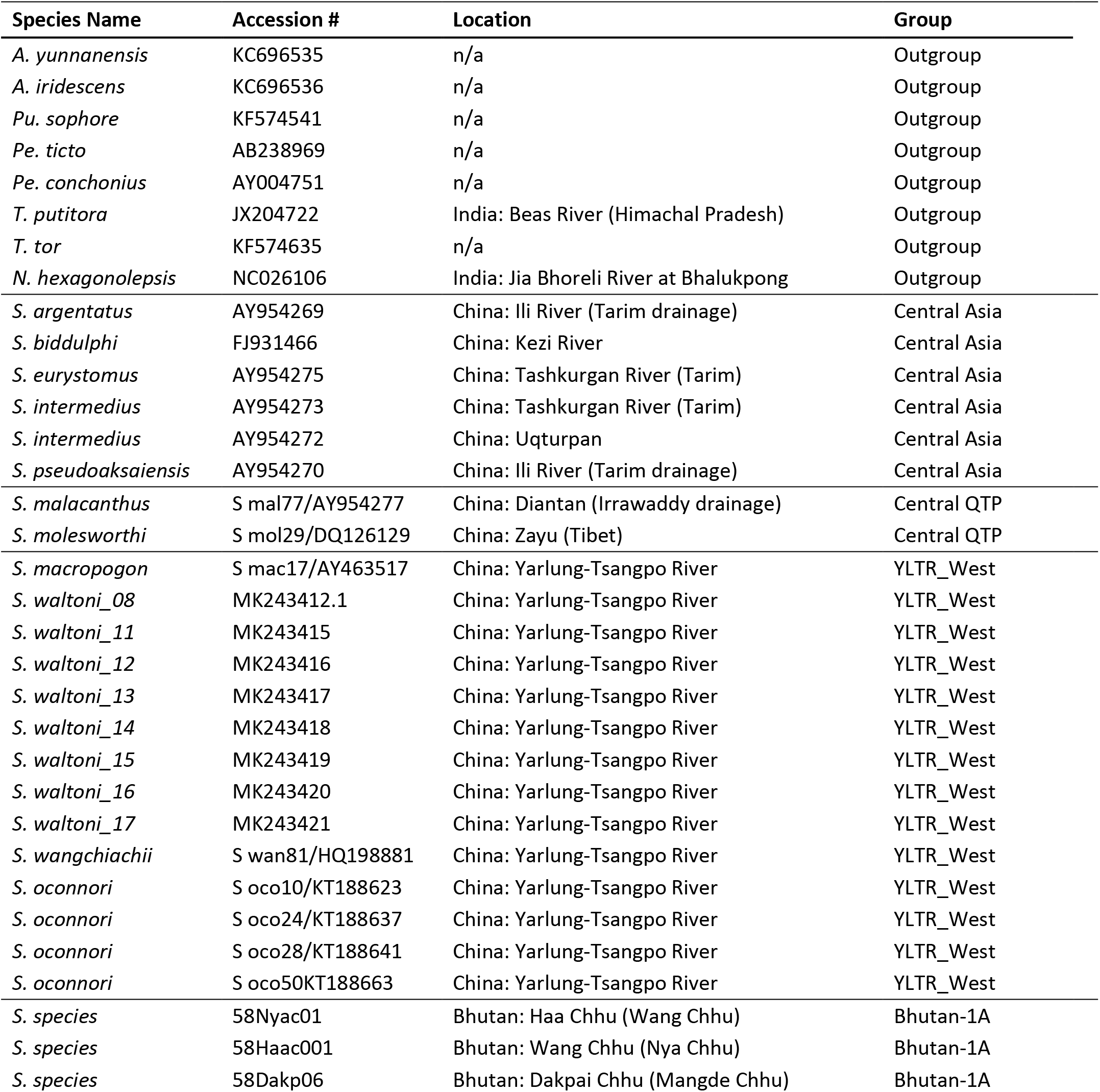
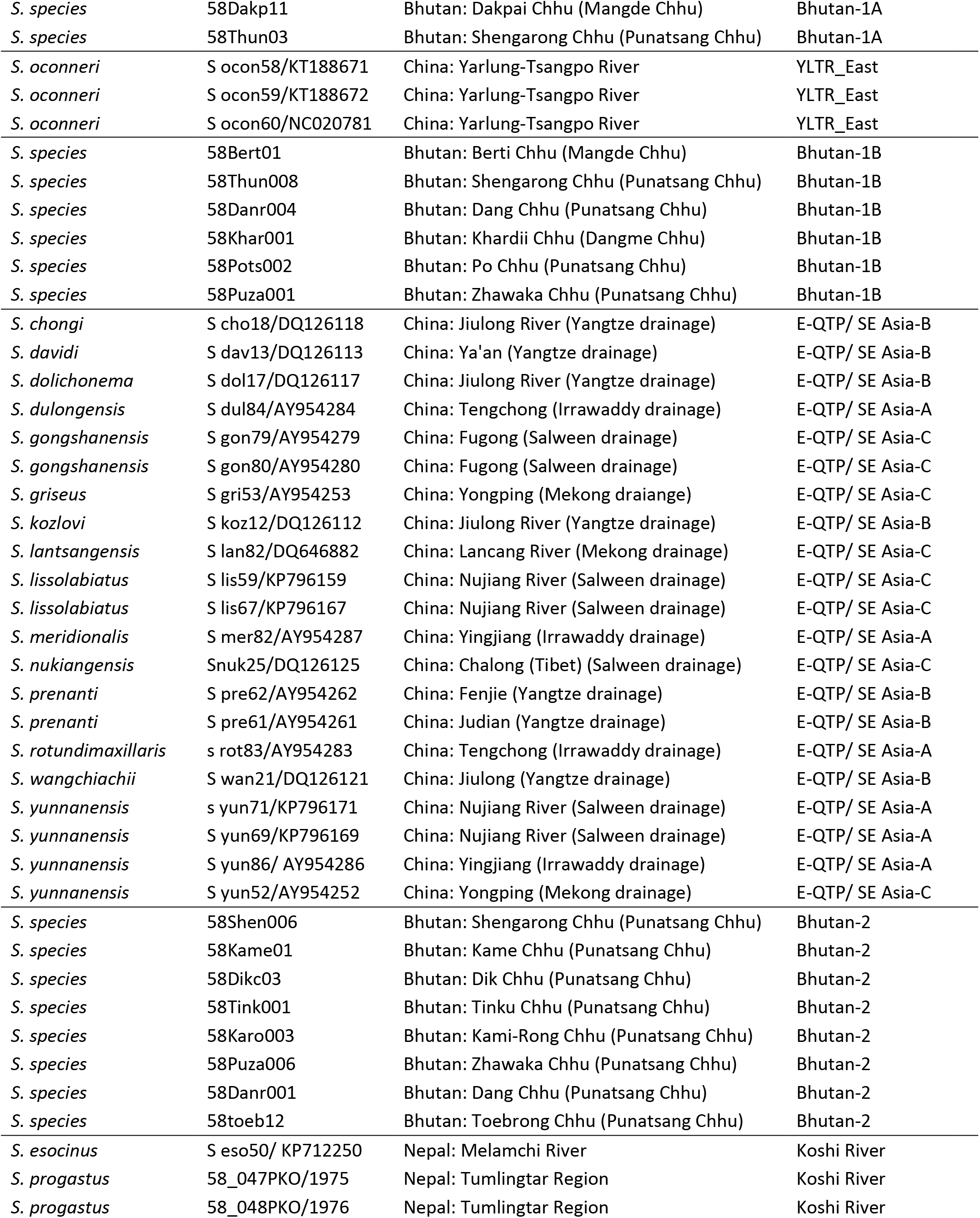
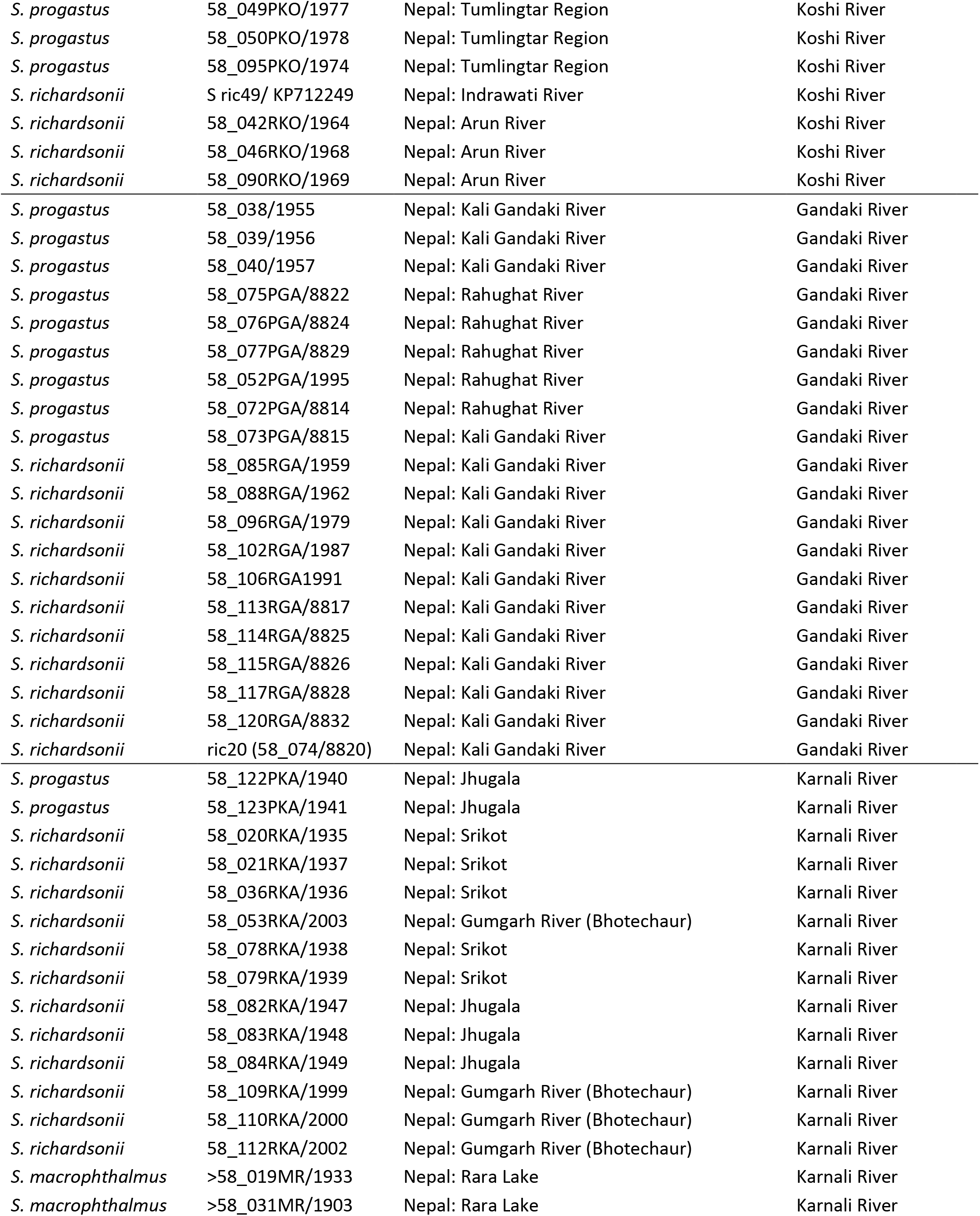
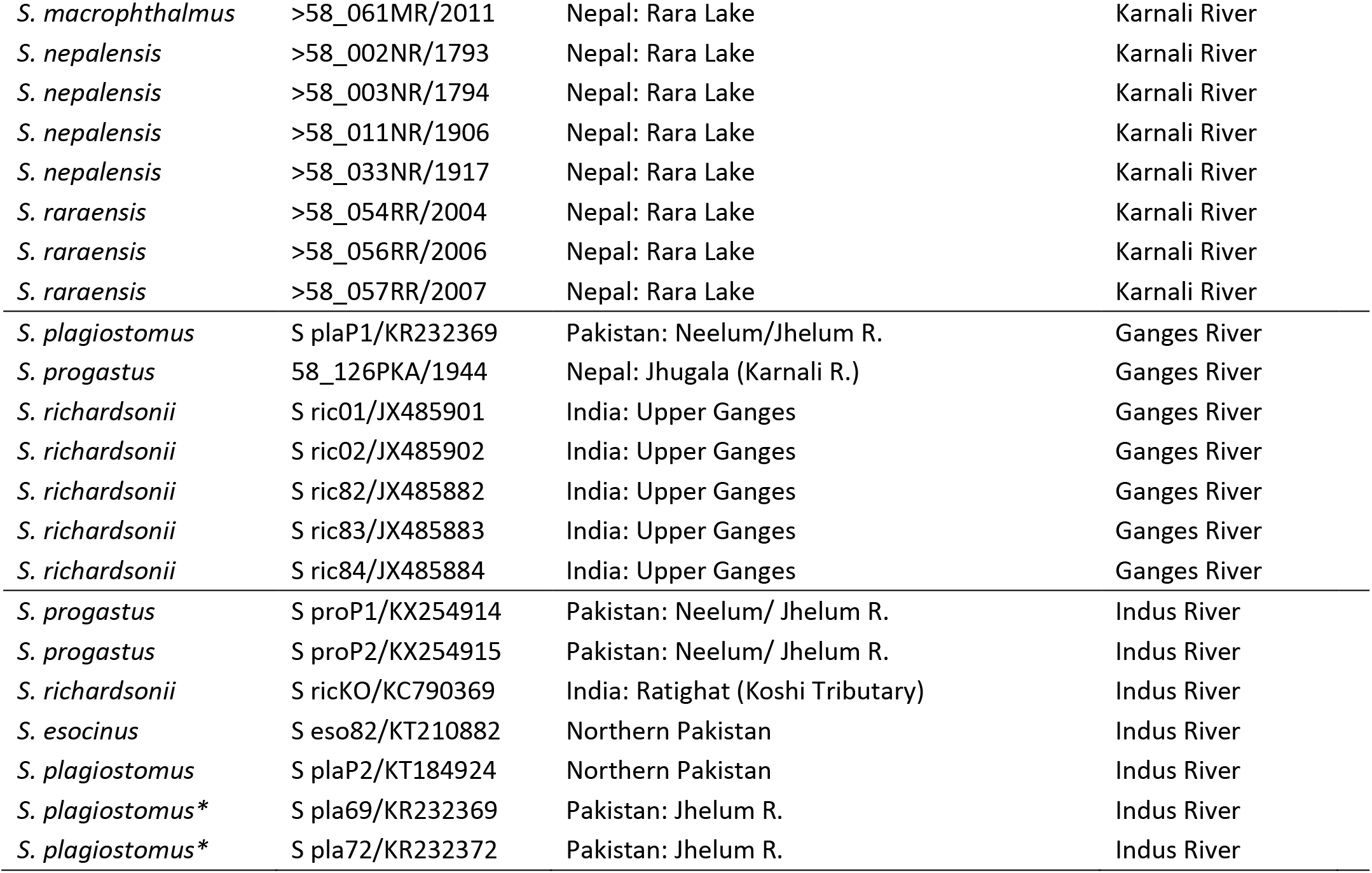
Fish biodiversity evaluated in this study. Listed are: Species Name = Scientific name (A=*Acrossocheilus*; Pe=*Pethia*; Pu=*Puntius*; T=*Tor*; N=*Neolissochilus*; S=*Schizothorax*) (* = Redundant haplotype); Accession# = Tissue or sequence identifier [i.e., GenBank accession number (begins with two letters); University of Kansas tissue voucher (four digits); Douglas Lab DNA code (begins with ‘S’ or ‘58’)]; Location = Country, geographic region or river; Group = Regions as presented in Figure 2; Group abbreviations are as follows: E-QTP/ SE Asia-A=Eastern Qinghai-Tibetan Plateau/ S.E. Asia, clade A; E-QTP/ SEA-B=Eastern QTP/S.E. Asia, clade B; E-QTP/ SEA-C=Eastern QTP/S.E. Asia, clade C; YLTR=Yarlung-Tsangpo River.

**Supplementary Figure S1:**
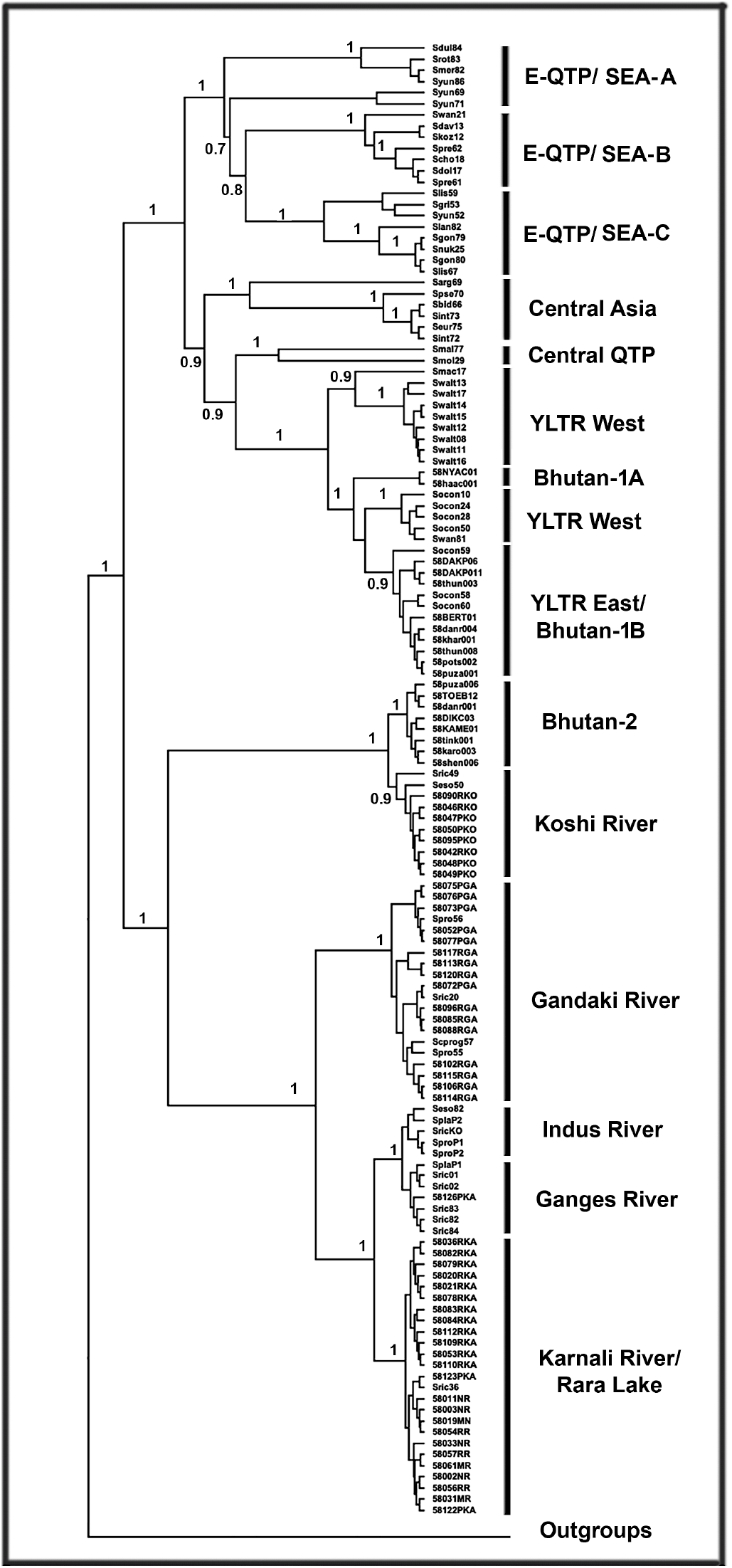
Phylogenetic tree derived using Bayesian methodology (BEAST v2.61) depicting phylogenetic relationships among Snowtrout (*Schizothora;* Cyprinidae) and outgroup taxa derived from sequence analysis of the cytochrome-b mitochondrial gene (1140 bp, 139 sequences, redundant haplotypes removed). Individuals are coalesced into geographic regions, with abbreviations defined as: E-QTP/ SEA-A (Eastern Qinghai-Tibetan Plateau/ S.E. Asia, clade A); E-QTP/ SEA-B (Eastern QTP/S.E. Asia, clade B); E-QTP/ SEA-C (Eastern QTP/S.E. Asia, clade C); YLTR (Yarlung-Tsangpo River). Numbers at nodes represent Bayesian posterior probabilities. Sample metadata in Table S1.

## Notes

### Competing Interest Statement

The authors have declared no competing interest.

### Summary of Updates

Title change and minor corrections

## References

[1] Yin, A. & Harrison, T. M. Geologic evolution of the Himalayan Tibetan orogen. Annu. Rev. Earth Planet. Sci., 28, 211–280. https://doi.org/10.1146/annurev.earth.28.1.211 (2000).

[2] Xu, J. & Grumbine, R. E. Building ecosystem resilience for climate change adaptation in the Asian highlands. WIREs Clim. Change, 5, 709–718. https://doi.org/10.1002/wcc.302 (2014).

[3] Pellissier, L., Heine, C., Rosauer, D. F. & Albouy, C. Are global hotspots of endemic richness shaped by plate tectonics? Biol. J. Linn. Soc., 123, 247–261. https://doi.org/10.1093/biolinnean/blx125 (2017).

[4] van Hinsbergen, D. J. et al. Greater India Basin hypothesis and a two-stage Cenozoic collision between India and Asia. Proc. Nat. Acad. Sci. U.S.A., 109, 7659–7664. https://doi.org/10.1073/pnas.1117262109 (2012).

[5] Brookfield, M. E. The evolution of the great river systems of southern Asia during the Cenozoic India-Asia collision: Rivers draining southwards. Geomorphology, 22, 285–312. https://doi.org/10.1016/S0169-555X(97)00082-2 (1998).

[6] He, D. & Chen, Y. Biogeography and molecular phylogeny of the genus *Schizothorax* (Teleostei: Cyprinidae) in China inferred from cytochrome b sequences. J. Biogeogr., 33, 1448–1460. https://doi.org/10.1111/j.1365-2699.2006.01510.x (2006).

[7] Bracciali, L., Najman, Y., Parrish, R. R., Akhter, S. H. & Millar, I. The Brahmaputra tale of tectonics and erosion: Early Miocene river capture in the Eastern Himalaya. Earth Planet. Sci. Lett., 415, 25–37. https://doi.org/10.1016/j.epsl.2015.01.022 (2015).

[8] Li, S. et al. The evolution of Yarlung Tsangpo River: Constraints from the age and provenance of the Gangdese conglomerates, southern Tibet. Gondwana Res., 41, 249–266. https://doi.org/10.1016/j.gr.2015.05.010 (2017).

[9] Govin, G. et al. Timing and mechanism of the rise of the Shillong Plateau in the Himalayan foreland. Geology, 46, 279–282. https://doi.org/10.1130/G39864.1 (2018).

[10] Favre, A. et al. The role of the uplift of the Qinghai-Tibetan Plateau for the evolution of Tibetan biotas. Biol. Rev., 90, 236–253. https://doi.org/10.1111/brv.12107 (2015).

[11] Deng, T. et al. Review: Implications of vertebrate fossils for paleo-elevations of the Tibetan Plateau. Glob. Planet. Change, 174, 58–69. https://doi.org/10.1016/j.gloplacha.2019.01.005 (2019).

[12] Schluter, D. Speciation, ecological opportunity, and latitude. Am. Nat., 187, 1–18. https://doi.org/10.1086/684193 (2016).

[13] Jacobsen, D., Milner, A. M., Brown, L. E. & Dangles, O. Biodiversity under threat in glacier-fed river systems. Nat. Clim. Change, 2, 361–364. https://doi.org/10.1038/NCLIMATE1435 (2012).

[14] Li, J. et al. Climate refugia of snow leopards in High Asia. Biol. Conserv. 203, 188–196. https://doi.org/10.1016/j.biocon.2016.09.026 (2016).

[15] Scherrer, D. & Körner, C. Topographically controlled thermal-habitat differentiation buffers alpine plant diversity against climate warming. J. Biogeogr., 38, 406–416. https://doi.org/10.1111/j.1365-2699.2010.02407.x (2011).

[16] Myers, N., Mittermeier, R. A., Mittermeier, C. G., da Fonseca, G. A. B. & Kent, J. Biodiversity hotspots for conservation priorities. Nature, 403, 853–858. https://doi.org/10.1038/35002501 (2000).

[17] Fricke, R., Eschmeyer, W. N. & van der Laan R. (eds). Eschmeyer’s catalog of fishes: Genera, species, references. http://researcharchive.calacademy.org/research/ichthyology/catalog/fishcatmain.asp (2020).

[18] Zhao, K. I. et al. The youngest split in sympatric Schizothoracine fish (Cyprinidae) is shaped by ecological adaptations in a Tibetan Plateau glacier lake. Mol. Ecol., 18, 3616–3628. https://doi.org/10.1111/j.1365-294X.2009.04274.x (2009).

[19] Yao, T. et al. Third pole environment (TPE). Environ. Dev., 3, 52–64. https://doi.org/10.1016/j.envdev.2012.04.002 (2012).

[20] IPCC (Intergovernmental Panel on Climate Change). IPCC Special Report on the Ocean and Cryosphere in a Changing Climate (eds. Pörtner, H.-O., Roberts, D.C., Masson-Delmotte, V., Zhai, P., Tignor, M., Poloczanska, E., Mintenbeck, K., Alegría, A., Nicolai, M., Okem, A., Petzold, J., Rama B. & Weyer, N.M.), Glan, Switzerland (2019).

[21] Zhu, G-F., Gao, L-M. & Li, D-Z. Protect third-pole’s fragile ecosystem. Science, 362, 1368. https://doi.org/10.1126/science.aaw0443 (2018).

[22] Sharma, A. Dubey, V.K., Johnson, J.A., Rawal, Y.K. & Sivakumara, K. Is there always space at the top? Ensemble modeling reveals climate-driven high-altitude squeeze for the vulnerable snow trout *Schizothorax richardsonii* in Himalaya. Ecol. Indic., 120, 106900. https://doi.org/10.1016/j.ecolind.2020.106900 (2020).

[23] Guillot, G., Mortier F. & Estoup, A. GENELAND: A computer package for landscape genetics. Mol. Ecol. Notes, 5, 712–715. https://doi.org/10.1111/j.1471-8286.2005.01031.x (2005).

[24] Guillot G. Inference of structure in subdivided populations at low levels of genetic differentiation—the correlated allele frequencies model revisited. Bioinformatics, 24, 2222–2228. https://doi.org/10.1093/bioinformatics/btn419 (2008).

[25] Excoffier, L. & Lischer H. E. L. ARLEQUIN suite ver 3.5: A new series of programs to perform population genetics analyses under Linux and Windows. Mol. Ecol. Resour., 10, 564–567. https://doi.org/10.1111/j.1755-0998.2010.02847.x (2010).

[26] Whipple, K. X. The influence of climate on the tectonic evolution of mountain belts. Nature Geosci., 2, 97–99. https://doi.org/10.1038/ngeo413 (2009).

[27] Hopken, M. W., Douglas, M. R. & Douglas, M. E. Stream hierarchy defines riverscape genetics of a North American desert fish. Mol. Ecol., 22, 956–971. https://doi.org/10.1111/mec.12156 (2013).

[28] Douglas, M. R., Brunner, P. C. & Douglas, M. E. Drought in an evolutionary context: Molecular variability in Flannelmouth Sucker (*Catostomus latipinnis*) from the Colorado River Basin of western North America. Freshw. Biol., 48, 1254–1273. https://doi.org/10.1046/j.1365-2427.2003.01088.x (2003).

[29] Douglas, M. R. et al. Anthropogenic impacts drive niche and conservation metrics of a cryptic rattlesnake on the Colorado Plateau of western North America. Royal Soc. Open Sci., 3, 160047. https://doi.org/10.1098/rsos.160047 (2016).

[30] Bishop, P. Drainage rearrangement by river capture, beheading and diversion. Prog. Phys. Geogr., 19, 449–473. https://doi.org/10.1177/030913339501900402 (1995).

[31] Strange, R. M. & Burr, B. M. Intraspecific phylogeography of North American Highland fishes: A test of the Pleistocene vicariance hypothesis. Evolution, 51, 885–897. https://doi.org/10.1111/j.1558-5646.1997.tb03670.x (1997).

[32] Bangs, M. R., Douglas, M. R., Mussmann, S. M. & Douglas, M. E. RUnraveling historical introgression and resolving phylogenetic discord within *Catostomus* (Osteichthys: Catostomidae). BMC Evol. Biol., 18, 86. https://doi.org/10.1186/s12862-018-1197-y (2018).

[33] Bangs, M. R., Douglas, M. R., Mussmann, S. M. & Douglas, M. E. Reticulate evolution as a management challenge: Patterns of admixture with phylogenetic distance in endemic fishes of Western North America. Evol. Applic., 13, 1400–1419. https://doi.org/10.1111/eva.13042 (2020).

[34] Chafin, T.K., Douglas, M.R. & Douglas M.E. Hybridization drives genetic erosion in sympatric desert fishes of western North America. Heredity, 123, 759–773. https://doi.org/10.1038/s41437-019-0259-2 (2019).

[35] Radinger, J. et al. The future distribution of river fish: The complex interplay of climate and land use changes, species dispersal and movement barriers. Glob. Change Biol., 23, 4970–4986. https://doi.org/10.1111/gcb.13760 (2017).

[36] CREP-Critical Ecosystem Partnership Fund. The biodiversity hotspots. http://www.cepf.net/resources/hotspots/Pages/default.aspx (2017).

[37] Qi, D. et al. The biogeography and phylogeny of schizothoracine fishes (*Schizopygopsis*) in the Qinghai-Tibetan Plateau. Zool. Scr., 44, 523–533. https://doi.org/10.1111/zsc.12116 (2015).

[38] Chen, W., Shen, Y., Gan, X., Wang, X. & He, S. Genetic diversity and evolutionary history of the *Schizothorax* species complex in the Lancang River (upper Mekong). Ecology and Evolution, 6, 6023–6036. https://doi.org/10.1002/ece3.2319 (2016).

[39] Chen, W., Yue, X. & He, S. Genetic differentiation of the *Schizothorax* species complex (Cyprinidae) in the Nujiang River (upper Salween). Sci. Rep., 7, 5944. https://doi.org/10.1038/s41598-017-06172-5 (2017).

[40] Wanghe, K. et al. Phylogeography of *Schizopygopsis stoliczkai* (Cyprinidae) in Northwest Tibetan Plateau area. Ecol. Evol., 7, 9602–9612. https://doi.org/10.1002/ece3.3452 (2017).

[41] Froese, R. & Pauly, D. (eds). FishBase 2000. World Wide Web electronic publication. www.fishbase.org (2019).

[42] Huo, T. B. et al. Length–weight relationships of 16 fish species from the Tarim River, China. J. Appl. Ichthyol., 28, 152–153. https://doi.org/10.1111/j.1439-0426.2011.01899.x (2012).

[43] Guo, X-Z. et al. Phylogeography of the threatened tetraploid fish, *Schizothorax waltoni*, in the Yarlung Tsangpo River on the southern Qinghai-Tibet Plateau: Implications for conservation. Sci. Rep., 9, 2704. https://doi.org/10.1038/s41598-019-39128-y (2019).

[44] Guo, X-Z. et al. Phylogeography and population genetics of *Schizothorax o’connori*: Strong subdivision in the Yarlung Tsangpo River inferred from mtDNA and microsatellite markers. Sci. Rep., 6, 29821. https://doi.org/10.1038/srep29821 (2016).

[45] Barat, A., Ali, S., Sati, J. & Sivaraman, G.K. Phylogenetic analysis of fishes of the subfamily Schizothoracinae (Teleostei: Cyprinidae) from Indian Himalayas using cytochrome b gene. Indian J. Fish., 59, 43–47. https://www.researchgate.net/publication/259772138 (2012)

[46] Terashima, A. Three new species of the cyprinid genus *Schizothorax* from Lake Rara, northwestern Nepal. Jap. J. Ichthyol., 31, 122–135. https://doi.org/10.11369/jji1950.31.122 (1984).

[47] Dimmick, W. W. & Edds, D. R. Evolutionary genetics of the endemic schizorathicine (Cypriniformes: Cyprinidae) fishes of Lake Rara, Nepal. Biochem. Syst. Ecol., 30, 919–929. https://doi.org/10.1016/S0305-1978(02)00030-3 (2002).

[48] Zakaria, G. Late Quaternary Seismicity and Climate in Western Nepal: Himalaya. Applied Geology. Unpubl. Ph.D. dissertation. Université Ghent, Department of Geology, Grenoble, France, Ghent, Belgium. https://tel.archives-ouvertes.fr/tel-01935412/document (2018).

[49] Martin, B.T. et al. Contrasting signatures of introgression in the contact zones of North American Box Turtle (*Terrapene* spp.). Mol. Ecol., 2020, 00:01–17. https://doi.org/10.1111/mec.15622 (2020).

[50] Mussmann, S.M., Douglas, M.R., Oakey, D.D. & Douglas M.E. Defining relictual biodiversity: Conservation units in Speckled Dace (Cyprinidae: *Rhinichthys osculus*) of the Greater Death Valley Ecosystem. Ecol. Evol., 2020, 00:1–20. https://doi.org/10.1002/ece3.6736) (2020).

[51] Qi, D. et al. Convergent, parallel and correlated evolution of trophic morphologies in the subfamily Schizothoracinae from the Qinghai-Tibetan Plateau. PLoS ONE, 7(3), e34070. https://doi.org/10.1371/journal.pone.0034070 (2012).

[52] Chirouze, F. et al. Detrital thermochronology and sediment petrology of the middle Siwaliks along the Muksar Khola section in eastern Nepal. J. Asian Earth Sci., 44, 94–106. https://doi.org/10.1016/j.jseaes.2011.01.009 (2013).

[53] Clift, P. D. & Blusztajn, J. Reorganization of the western Himalayan river system after five million years. Nature, 438, 1001–1003. https://doi.org/10.1038/nature04379 (2013).

[54] Braulik, G. T. et al. One species or two? Vicariance, lineage divergence and low mtDNA diversity in geographically isolated populations of South Asian River dolphin. J. Mammal. Evol., 22, 111–120. https://doi.org/10.1007/s10914-014-9265-6 (2015).

[55] Tang, Y. et al. Convergent evolution misled taxonomy in Schizothoracine fishes (Cypriniformes: Cyprinidae). Mol. Phylogenet. Evol., 134, 323–337. https://doi.org/10.1016/j.ympev.2019.01.008 (2019).

[56] Vilgalys, R. Taxonomic misidentification in public DNA databases. New Phytol., 160, 4–5. https://doi.org/10.1046/j.1469-8137.2003.00894.x (2003).

[57] Edds, D. R. Fishes in Nepal: Ichthyofaunal surveys in seven nature reserves. Ichthyol. Explor. Freshw., 18, 277–287. https://www.researchgate.net/publication/228787571_Fishes_in_Nepal_Ichthyofaunal_surveys_in_seven_nature_reserves (2007).

[58] Yang, J., Yang, J. X. & Chen, X. Y. A re-examination of the molecular phylogeny and biogeography of the genus *Schizothorax* (Teleostei: Cyprinidae) through enhanced sampling with an emphasis on the species in the Yunnan-Guizhou plateau, China. J. Zool. Syst. Evol. Res., 50, 184–191. https://doi.org/10.1111/j.1439-0469.2012.00661.x (2012).

[59] Sharma, C. M. Freshwater fishes, fisheries, and habitat prospects of Nepal. Aquat. Ecosys. Health Manag., 11, 289–297. https://doi.org/10.1080/14634980802317329 (2008).

[60] Das, S. M. & Subla, B. A. The ichthyofauna of Kasmir: Part II The speciation of Kasmir fishes with two new records of species. Ichthyologica, 3, 57–62 (1964).

[61] Cao, W. X., Chen, Y. Y., Wu, Y. F. & Zhu, S. Q. Origin and evolution of Schizothoracine fishes in relation to the upheaval of the Qinghai-Xizang Plateau in Studies on the Period, Amplitude and Type of the Uplift of the Qinghai-Xizang Plateau (ed. Li, Q. F.) 118–129 (Beijing, China: Science Press, 1981).

[62] Yang, L. et al. Phylogeny and polyploidy: Resolving the classification of cyprinine fishes (Teleostei: Cypriniformes). Mol. Phylogenet. Evol., 85, 97–116. https://doi.org/10.1016/j.ympev.2015.01.014 (2015).

[63] Wang, X., Gan, X., Li, Y. & He, S. Cyprininae phylogeny revealed independent origins of the Tibetan Plateau endemic polyploid cyprinids and their diversifications related to the Neogene uplift of the plateau. Sci. China Life Sci., 59, 1149–1165. https://doi.org/10.1007/s11427-016-0007-7 (2016).

[64] Taylor, J.S., Braasch, I., Frickey, T., Meyer, A. & Van de Peer, Y. Genome duplication, a trait shared by 22,000 species of ray-finned fish. Genome Res., 13, 382–390. https://doi.org/10.1101/gr.640303 (2003).

[65] Groth, J. G. & Barrowclough, G. F. Basal divergences in birds and the phylogenetic utility of the nuclear RAG-1 gene. Mol. Phylogenet. Evol., 12, 115–123. https://doi.org/10.1006/mpev.1998.0603 (1999).

[66] Houston, D. D., Evans, R. P. & Shiozawa, D. K. Evaluating the genetic status of a Great Basin endemic minnow: The relict dace (*Relictus solitarius*). Cons. Gen., 13, 727–742. https://doi.org/10.1007/s10592-012-0321-6 (2012).

[67] Kumar, S. Stecher, G., Li, M., Knyaz, C. & Tamura, K., MEGA X: Molecular Evolutionary Genetics Analysis across computing platforms. Mol. Biol. Evol., 35, 1547–1549. https://doi.org/10.1093/molbev/msy096 (2018).

[68] Rozas, J. et al. DNASP 6: DNA sequence polymorphism analysis of large datasets. Mol. Biol. Evol. 34, 3299–3302. https://doi.org/10.1093/molbev/msx248 (2017).

[69] R-Studio Team. RStudio: Integrated Development for R. RStudio, PBC, Boston, MA. http://www.rstudio.com/ (2020).

[70] Solecki, A. M., Skevington, J. H., Buddle, C. M. & Wheeler, T. A. Phylogeography of higher Diptera in glacial and postglacial grasslands in western North America. BMC Ecol., 19, 53. https://doi.org/10.1186/s12898-019-0266-4 (2019).

[71] Minh, B. Q. et al. IQ-TREE 2: New models and efficient methods for phylogenetic inference in the genomic era. Mol. Biol. Evol., 37, 1530–1534. https://doi.org/10.1093/molbev/msaa015 (2020).

[72] Kalyaanamoorthy, S., Minh, B. Q., Wong, T. K. F., von Haeseler, A. & Jermiin, L. S. MODELFINDER: Fast model selection for accurate phylogenetic estimates Nat. Methods, 14, 587–589. https://doi.org/10.1038/nmeth.4285 (2017).

[73] Hoang, D. T., Chernomor, O., von Haeseler, A., Minh, B. Q. & Vinh, L. S. UFBOOT2: Improving the ultrafast bootstrap approximation. Mol. Biol. Evol., 35, 518–522. https://doi.org/10.1093/molbev/msx281 (2018).

[74] Drummond, A. J., Suchard, M. A., Xie, D. & Rambaut, A. Bayesian phylogenetics with BEAUTI and the BEAST 1.7. Mol. Biol. Evol., 29, 1969–1973. https://doi.org/10.1093/molbev/mss075 (2012).

[75] Rambaut, A. & Drummond, A. J. TRACER V1.4. Molecular evolution, phylogenetics and epidemiology. Edinburgh: The University of Edinburgh, Institute of Evolutionary Biology software. http://tree.bio.ed.ac.uk/software/tracer/ (2007).

[76] Rambaut, A. FIGTREE v1.4. Molecular evolution, phylogenetics and epidemiology. Edinburgh: The University of Edinburgh, Institute of Evolutionary Biology software. http://tree.bio.ed.ac.uk/software/figtree/ (2012).

[77] Kishino, H., Miyata, T. & Hasegawa, M. Maximum likelihood inference of protein phylogeny and the origin of chloroplasts. J. Mol. Evol., 31, 151–160. https://doi.org/10.1007/BF02109483 (1990).

[78] Kishino, H. & Hasegawa, M. Evaluation of the maximum likelihood estimate of the evolutionary tree topologies from DNA sequence data, and the branching order in hominoidea. J. Mol. Evol., 29, 170–179. https://doi.org/10.1007/BF02100115 (1989).

[79] Shimodaira, H. & Hasegawa, M. Multiple comparisons of log-likelihoods with applications to phylogenetic inference. Mol. Biol. Evol., 16, 1114–1116. https://doi.org/10.1093/oxfordjournals.molbev.a026201 (1999).

[80] Shimodaira, H. An approximately unbiased test of phylogenetic tree selection. Syst. Biol., 51, 492–508. https://doi.org/10.1080/10635150290069913 (2002).

[81] Strimmer, K. & Rambaut A. Inferring confidence sets of possibly misspecified gene trees. Proc. R. Soc. London. Ser. B Biol. Sci., 269, 137–142. https://doi.org/10.1098/rspb.2001.1862 (2002).

[82] Tamura, K., Tao, Q. & Kumar, S. Theoretical foundation of the RELTIME method for estimating divergence times from variable evolutionary rates. Mol. Biol. Evol., 35, 1770–1782. https://doi.org/10.1093/molbev/msy044 (2018).

[83] Mello, B. Estimating timetrees with MEGA and the TimeTree Resource. Mol. Biol. Evol., 35, 2334–2342. https://doi.org/10.1093/molbev/msy133 (2018).

[84] Yang, T. et al. New schizothoracine from Oligocene of Qaidam Basin, northern Tibetan Plateau, China, and its significance. J. Vertebr. Paleontol., 2017, e1442840. https://doi.org/10.1080/02724634.2018.1442840 (2018).

[85] Guo, J. et al. Three-dimensional structural model of the Qaidam Basin: Implications for crustal shortening and growth of the northeast Tibet. Open Geosci., 9,174–185. https://doi.org/10.1515/geo-2017-0015 (2017).

[86] Loucks, D.P. & van Beek, E. Water Resource Systems Planning and Management, An Introduction to Methods, Models, and Applications. Deltares, UNESCO-IHE, and Springer-Verlag Open Access. https://doi.org/10.1007/978-3-319-44234-1_1 (Fig. 1.17 China, India, and Southeast Asia, highlighting the Tibetan Plateau: Available via license: CC BY-NC 4.0) (2017).

